# Persistence of regenerative stem cells provides a cellular basis for the parasitostatic activity of albendazole against *Echinococcus multilocularis*

**DOI:** 10.64898/2026.09.11.750979

**Authors:** Monika Bergmann, Markus Spiliotis, Uriel Koziol, Klaus Brehm

## Abstract

Albendazole (ABZ) is the mainstay of chemotherapy for alveolar echinococcosis caused by the larval stage of *Echinococcus multilocularis*, but its activity is predominantly parasitostatic rather than parasiticidal. The cellular basis of this remarkable parasite persistence remains poorly understood. Here, we investigated the effects of sustained ABZ exposure on the germinative stem cells that drive metacestode growth and regeneration and examined whether differential β-tubulin isoform usage could contribute to their drug susceptibility. ABZ rapidly impaired microtubule-dependent tegumental transport and caused pronounced damage to metacestode vesicles. In contrast, proliferative germinative cells (GC) persisted during prolonged and continuous ABZ exposure. Importantly, cells surviving 14–28 days of treatment retained developmental competence, regenerated metacestode vesicles following drug withdrawal, and, after extended exposure to 10 µM ABZ, remained capable of generating parasite tissue following transplantation into a permissive host. Transcriptomic and immunohistochemical analyses identified the Y200-containing β-tubulin isoform Tub-2 as a major metacestode β-tubulin that is almost exclusively associated with the proliferative GC lineage. Tub-2 was present in virtually all S-phase cells, colocalized with established GC markers, localized to mitotic spindles, and Tub-2+ cells persisted during prolonged ABZ and ABZ-SO exposure. Structural analysis further revealed an E198–Y200 interaction in Tub-2, providing a structural correlate of the resistance-associated Y200 configuration. Together, these findings provide a cellular explanation for the predominantly parasitostatic activity of benzimidazoles against *E. multilocularis*. ABZ profoundly damages parasite tissue while sparing a regenerative cell compartment capable of re-establishing metacestode growth after drug withdrawal. They further identify GC-associated Tub-2 as a plausible molecular contributor to this persistence and as a potential target for curative anti-*Echinococcus* chemotherapy.

**Author Summary:** Alveolar echinococcosis is a severe parasitic disease caused by the larval stage of *E. multilocularis*. Treatment relies mainly on albendazole (ABZ), which can control parasite growth but usually does not eliminate the infection, making long-term or even lifelong therapy necessary. The biological reasons for this remarkable persistence are poorly understood. Here, we show that ABZ severely damages parasite tissue while a population of proliferating germinative stem cells survives even prolonged drug exposure. Importantly, these cells retain the ability to regenerate parasite tissue after ABZ is removed. We further identify the β-tubulin isoform Tub-2 as being strongly associated with these germinative cells and show structural features that may contribute to reduced ABZ susceptibility. Our findings suggest that the failure of ABZ to eliminate the parasite may result from its inability to eradicate the regenerative stem cell compartment. Drugs that specifically eliminate these cells could therefore provide a route towards curative treatment of alveolar echinococcosis.

## Introduction

Alveolar echinococcosis (AE), caused by the larval stage of the fox tapeworm *E. multilocularis*, is a severe and potentially lethal parasitic disease characterized by infiltrative growth of the metacestode, predominantly within the liver. Radical surgical resection can be curative, but is feasible only in a proportion of patients. For unresectable disease, treatment relies almost exclusively on the benzimidazole derivatives albendazole (ABZ) and mebendazole (MBZ), which have remained the mainstay of chemotherapy for several decades (Brunetti et al., 2010; Autier et al., 2024; Deibel et al., 2025). Although ABZ/MBZ treatment has dramatically improved the prognosis of AE, its activity is considered predominantly parasitostatic rather than parasiticidal. Consequently, treatment must usually be continued for many years and, in patients with unresectable disease, is frequently lifelong (Brunetti et al., 2010; Grüner et al., 2017; Deibel et al., 2025). The capacity of parasite tissue to resume growth after discontinuation of therapy has been documented clinically, including recurrence after several years of continuous benzimidazole treatment (Ammann et al., 1990). Although parasite inactivation permitting treatment discontinuation can occur in a subset of patients following long-term therapy, reliable demonstration of parasite death remains difficult and recurrence may still occur after treatment cessation (Ammann et al., 2015; Deibel et al., 2022). Thus, there is currently no reliably curative chemotherapy for AE (Autier et al., 2024). The cellular basis underlying this remarkable persistence of *E. multilocularis* during benzimidazole treatment remains poorly understood.

A distinctive feature of the *E. multilocularis* metacestode is its population of germinative stem cells. Germinative cells (GC) constitute the only continuously proliferating somatic cell population of the larval parasite, account for 20–25% of all metacestode cells, and are responsible for generating the differentiated cell types required for tissue homeostasis, growth, and development (Koziol et al., 2014; Herz et al., 2024). They are further characterized by a remarkable regenerative potential: primary cell preparations enriched in GC can regenerate complete metacestode vesicles *in vitro* and give rise to parasite tissue following transplantation into suitable hosts (Spiliotis et al., 2008; Koziol et al., 2014). Accordingly, pharmacological depletion or disruption of the GC compartment profoundly impairs parasite development, highlighting these cells as an important target for curative chemotherapy (Schubert et al., 2014; Cheng et al., 2019). Conversely, damage to differentiated parasite tissues would not necessarily result in parasite death if developmentally competent GC persist. Based on these considerations, we previously proposed that the relative refractoriness of GC to benzimidazoles might contribute to the predominantly parasitostatic activity of these drugs and provide a cellular reservoir from which parasite growth can resume following treatment cessation (Brehm and Koziol, 2014; Koziol and Brehm, 2015). However, whether GC indeed survive prolonged and continuous benzimidazole exposure while retaining proliferative and regenerative capacity has not been directly established.

The primary molecular target of benzimidazoles is β-tubulin, and binding of these compounds interferes with microtubule assembly and consequently with microtubule-dependent processes such as intracellular transport and cell division (Lacey and Gill, 1994). Extensive studies in parasitic nematodes have established variation in β-tubulin sequence and isoform expression as major determinants of benzimidazole susceptibility. In particular, substitutions at residues 167, 198, and 200 have repeatedly been associated with benzimidazole resistance, with F200Y representing one of the most extensively characterized resistance-associated substitutions (Kwa et al., 1994; von Samson-Himmelstjerna et al., 2007; Furtado et al., 2014; Dilks et al., 2020). More recently, introduction of F167Y, E198A, or F200Y into a benzimidazole-susceptible *Caenorhabditis elegans* background provided direct functional evidence that each of these substitutions is sufficient to confer resistance (Dilks et al., 2020). Structural modelling and molecular dynamics studies further indicate that these residues contribute to the local architecture of the benzimidazole-binding region, with E198 participating directly in drug interactions and substitutions at position 200 affecting the interaction network surrounding this residue (Aguayo-Ortiz et al., 2013a, b). *E. multilocularis* expresses multiple β-tubulin isoforms with distinct sequence characteristics, including isoforms containing either phenylalanine or tyrosine at position 200 (Brehm et al., 2000; Tsai et al., 2013). Previous transcriptomic observations suggested that the Y200-containing isoform Tub-2 might be preferentially expressed under conditions enriched in GC (Herz et al., 2024), leading us to propose that expression of a β-tubulin isoform with reduced benzimidazole susceptibility could contribute to the persistence of the parasite stem cell compartment during chemotherapy (Schubert et al., 2014; Brehm and Koziol, 2014; Koziol and Brehm, 2015). However, this hypothesis has remained untested: the cellular distribution and function of Tub-2 within the metacestode have not been established, nor has it been determined whether Tub-2-expressing cells preferentially persist during prolonged benzimidazole exposure.

Here, we directly tested whether the GC compartment of *E. multilocularis* persists during prolonged ABZ exposure and investigated whether β-tubulin isoform usage could provide a molecular correlate of this differential drug susceptibility. Using sustained drug exposure of intact metacestodes and GC-enriched primary cultures, we show that ABZ profoundly damages metacestode vesicles and rapidly disrupts microtubule-dependent tegumental transport, while proliferative and developmentally competent GC persist even after several weeks of treatment. We further identify the Y200-containing β-tubulin isoform Tub-2 as a major metacestode β-tubulin that is preferentially associated with the proliferative GC compartment.

Structural analysis revealed an E198–Y200 interaction in Tub-2, providing a structural correlate of the resistance-associated Y200 configuration, while Tub-2+ cells persisted during prolonged ABZ and ABZ-SO exposure. Together, these findings provide a cellular basis for the predominantly parasitostatic activity of benzimidazoles against *E. multilocularis* and identify GC-associated Tub-2 as a potential molecular contributor to the relative refractoriness of the parasite stem cell compartment.

## Materials and Methods

### Parasite material and *in vitro* cultivation

*E. multilocularis* metacestode material of isolates H95 and GH09 was maintained by serial intraperitoneal passage in *Meriones unguiculatus* and subsequently cultivated under axenic conditions as previously described (Spiliotis et al., 2008; Spiliotis and Brehm, 2009). Metacestode vesicles were maintained in conditioned culture medium under reducing conditions at 37°C and 5% CO₂, with medium changes every three days. Primary cell cultures were established from axenically cultivated metacestode vesicles according to the previously established procedure (Spiliotis et al., 2008; Spiliotis and Brehm, 2009). Briefly, parasite tissue was mechanically and enzymatically dissociated, and isolated cells were maintained under conditions supporting aggregate formation and subsequent regeneration of metacestode vesicles. Unless stated otherwise, experiments were performed with isolate H95. For the long-term EdU (5-ethynyl-2′-deoxyuridine) experiments shown in Fig. 2, three independent experiments were performed using two preparations of isolate GH09 and one preparation of H95. No apparent isolate-dependent differences were observed. Experiments shown in Figs. 6 and 7 were performed with isolate GH09. Protoscoleces used for Fig. 12 were derived from isolate GH09; all other experiments shown in Figs. 10–14 were performed with H95. Protoscoleces were isolated and activated as previously described (Herz et al., 2024).

**Figure 1.**
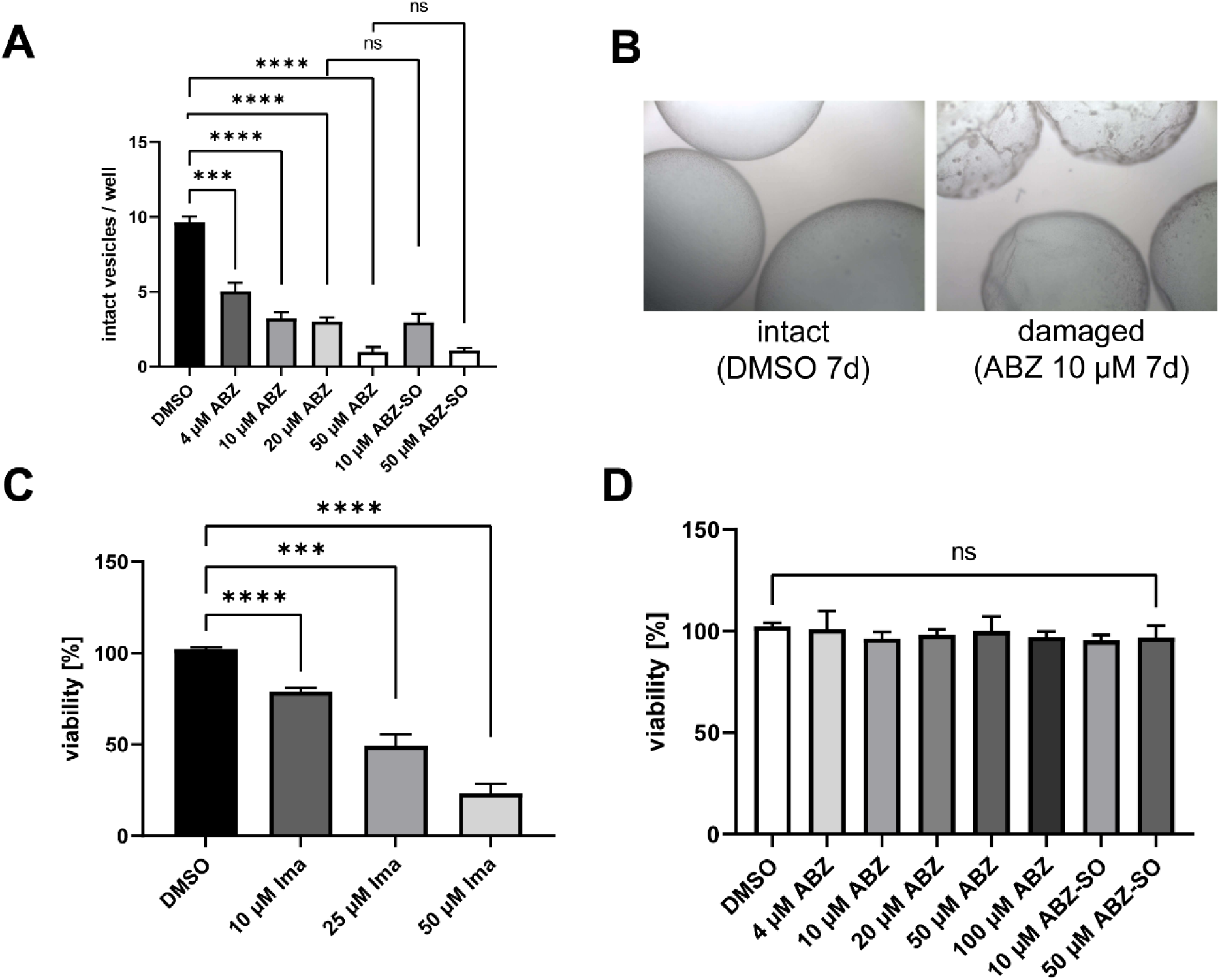
ABZ disrupts metacestode vesicles but does not reduce the viability of stem cell-enriched primary cells. **(A)** Effect of ABZ and ABZ-SO on *Echinococcus multilocularis* metacestode vesicles. Vesicles were cultured in the presence of the indicated concentrations of ABZ or ABZ-SO for 7 days, and the number of intact vesicles per well was determined. DMSO-treated cultures served as controls. **(B)** Representative images illustrating the morphological criteria used to distinguish intact and damaged metacestode vesicles. Shown are intact vesicles after 7 days of DMSO treatment and damaged vesicles after treatment with 10 µM ABZ for 7 days. **(C)** Effect of imatinib (Ima) on the viability of *E. multilocularis* primary cells as determined by resazurin reduction assay. Primary cells were treated with the indicated concentrations of imatinib, and viability was expressed relative to the DMSO-treated control, which was set to 100%. **(D)** Effect of ABZ and ABZ-SO on the viability of *E. multilocularis* primary cells as determined by resazurin reduction assay. Primary cells were treated with the indicated concentrations of ABZ or ABZ-SO, and viability was expressed relative to the DMSO-treated control, which was set to 100%. For all experiments, three independent biological replicates were performed, each comprising three technical replicates. Technical replicates were averaged within each biological replicate, and these biological replicate means were used as individual values for statistical analysis. Bars represent the mean ± SD of the three biological replicates. Statistical significance relative to the corresponding DMSO control was assessed using unpaired two-tailed Student’s *t*-tests. ns, not significant; \*\*\**P* < 0.001; \*\*\*\**P* < 0.0001.

**Figure 2.**
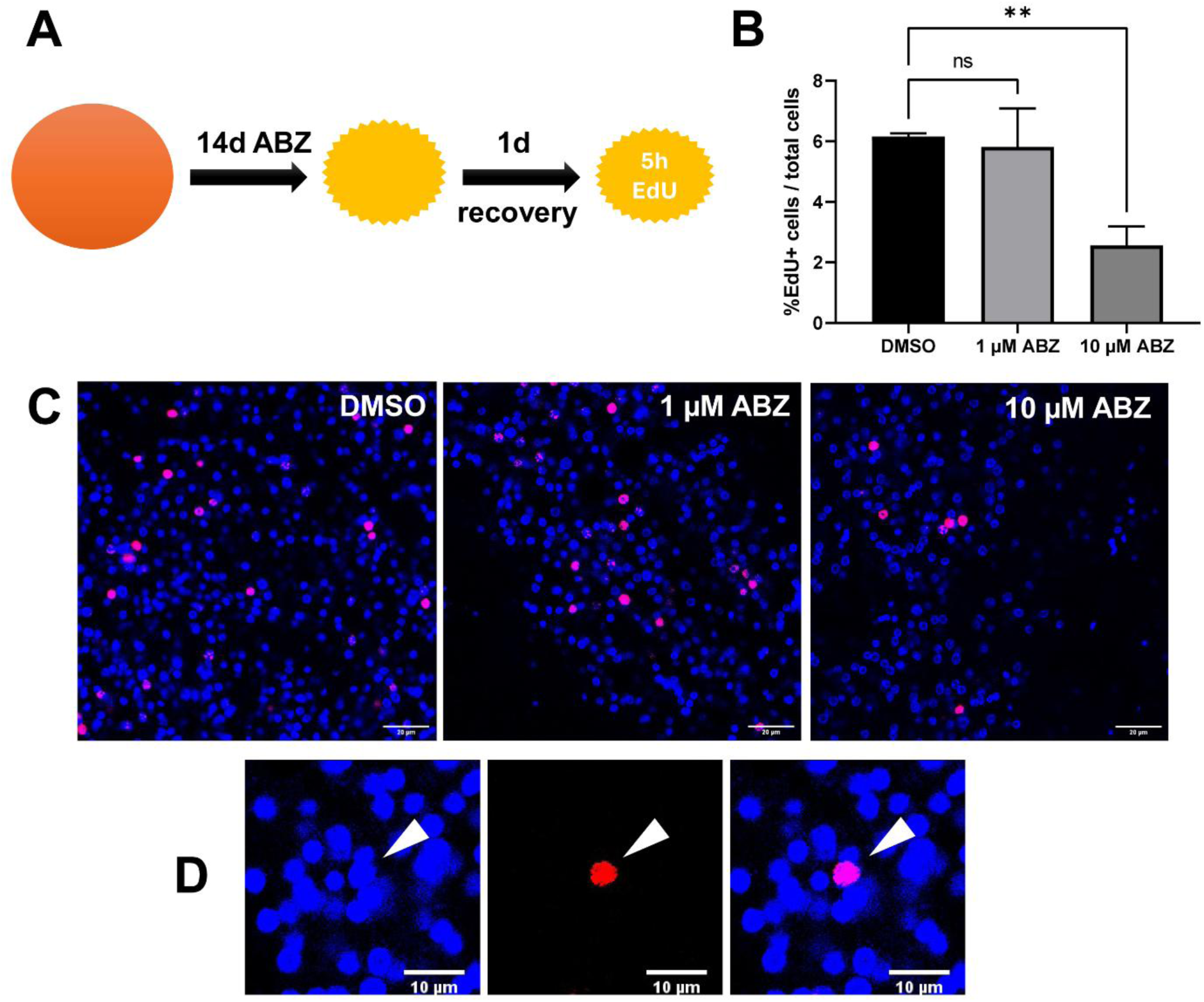
Proliferative stem cells persist after prolonged albendazole treatment of metacestode vesicles. **(A)** Experimental setup. *E. multilocularis* metacestode vesicles were treated with DMSO or the indicated concentrations of ABZ for 14 days, followed by a 1-day drug-free recovery period and a 5-h EdU pulse to label proliferating cells. **(B)** Quantification of EdU+ cells after ABZ treatment. The percentage of EdU+ cells relative to the total number of cells was determined for DMSO-treated controls and vesicles treated with 1 or 10 µM ABZ. Three independent biological replicates were analyzed. For each biological replicate, three technical replicates were included, with three vesicles analyzed per technical replicate. Three randomly selected microscopic fields were quantified per vesicle. The mean calculated from all quantified fields within each biological replicate was used as a single value for statistical analysis. Bars represent the mean ± SD of the three biological replicates. Statistical significance relative to the DMSO control was assessed using unpaired two-tailed Student’s *t*-tests. ns, not significant; \*\**P* < 0.01. **(C)** Representative fluorescence images of EdU labeling in metacestode vesicles treated with DMSO, 1 µM ABZ, or 10 µM ABZ. EdU+ nuclei are shown in magenta and total nuclei in blue. Scale bar, 20 µm. **(D)** Higher-magnification view of an EdU+ cell illustrating the nuclear localization of the EdU signal (arrowhead). Shown are the nuclear counterstain (blue), EdU signal (red), and merged image. Scale bar, 10 µm.

**Figure 3.**
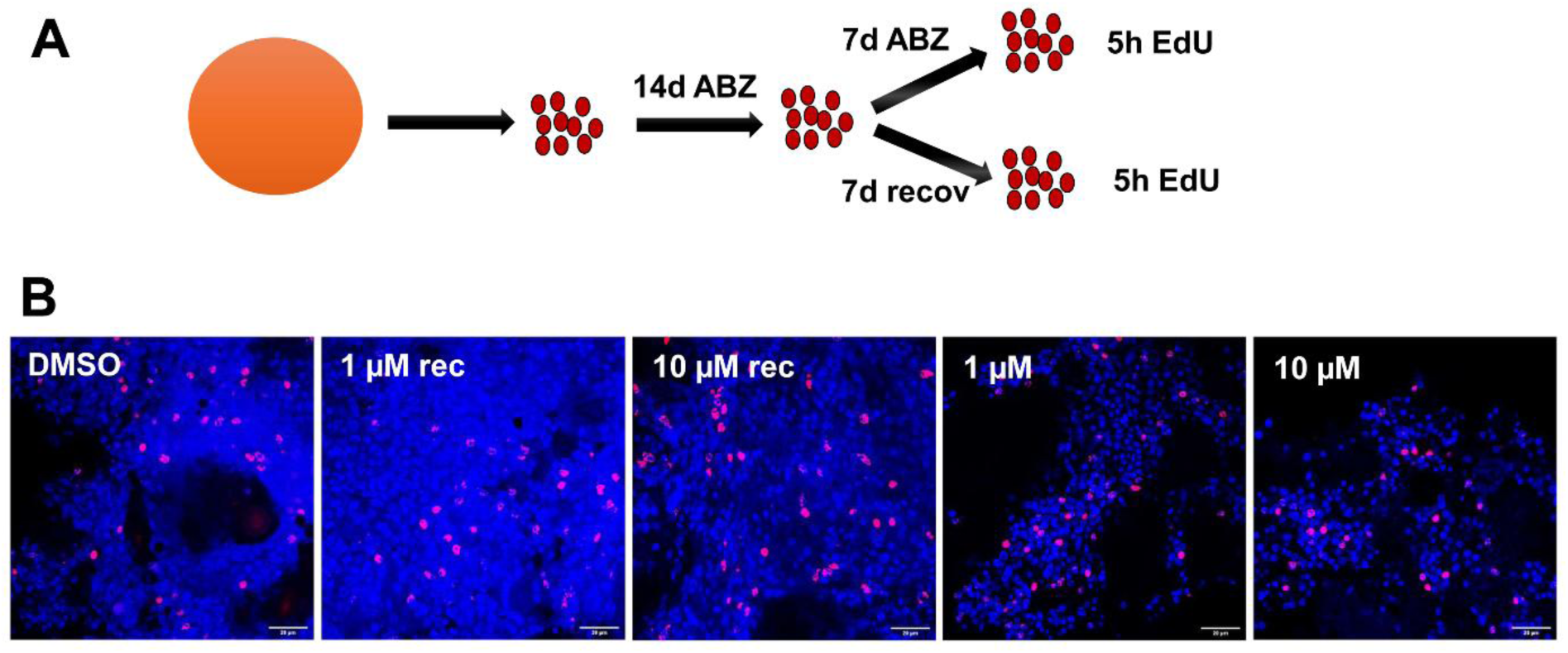
Proliferative cells persist in primary cell cultures during prolonged ABZ exposure. **(A)** Experimental setup. Primary cells were isolated from untreated *E. multilocularis* metacestode vesicles and cultured in the presence of DMSO, 1 µM ABZ, or 10 µM ABZ for 14 days. Cultures were subsequently either maintained for an additional 7 days in the presence of the respective ABZ concentration or subjected to a 7-day drug-free recovery period. At the end of the experiment, cells were exposed to a 5-h EdU pulse to label cells undergoing DNA synthesis. **(B)** Representative fluorescence images of primary cell aggregates at the end of the experiment. EdU+ nuclei are shown in magenta and total nuclei in blue. Conditions comprise DMSO-treated controls, cells treated with 1 or 10 µM ABZ for 14 days followed by 7 days of drug-free recovery (rec), and cells continuously exposed to 1 or 10 µM ABZ for 21 days. EdU-positive cells remained detectable after prolonged ABZ treatment, including after 21 days of continuous exposure without a drug-free recovery period. Images are representative of the observed primary cell aggregates; no quantitative comparison was performed because reliable quantification of EdU+ cells within the heterogeneous three-dimensional primary cell aggregates was not feasible. Scale bar, 20 µm.

**Figure 4.**
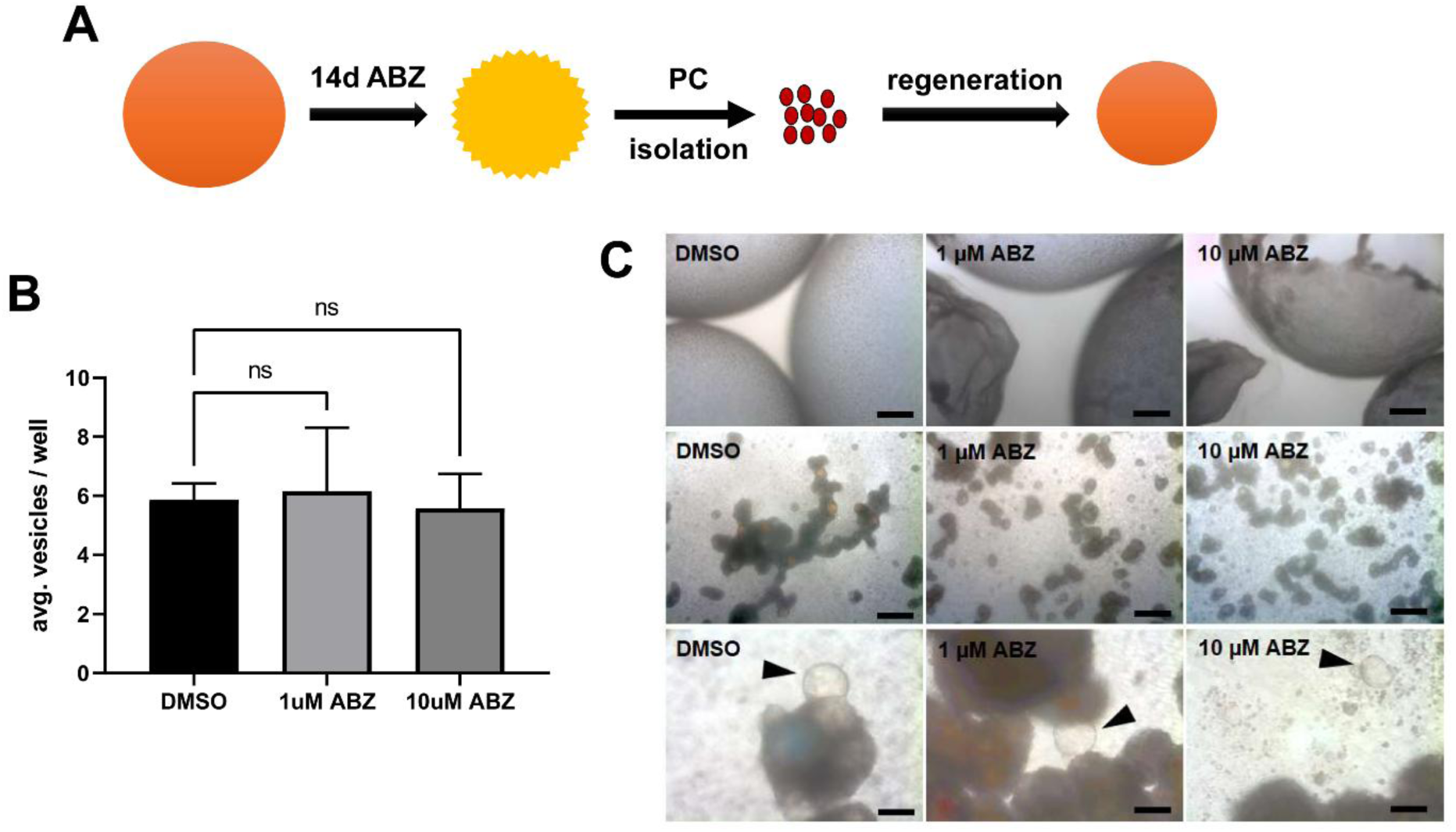
ABZ treatment does not impair the regenerative capacity of metacestode-derived primary cells. **(A)** Experimental design. Intact *E. multilocularis* metacestode vesicles were treated with DMSO (control), 1 µM ABZ, or 10 µM ABZ for 14 days. Primary cells were subsequently isolated from the treated vesicles and cultured for an additional 14 days in drug-free medium to assess their capacity to regenerate metacestode vesicles. **(B)** Number of newly formed metacestode vesicles after 14 days of drug-free primary cell culture. Three independently prepared metacestode vesicle cultures were treated and subsequently used in their entirety for primary cell isolation, yielding three independent biological replicates per condition without technical replicates. Each primary cell preparation was cultured separately, and newly formed vesicles were counted after 14 days. Data are shown as mean ± SD (*n* = 3 biological replicates). Statistical comparisons were performed using t-test. ns, not significant. **(C)** Representative images illustrating the experiment. Upper panels show metacestode vesicles after 14 days of treatment with DMSO, 1 µM ABZ, or 10 µM ABZ. Middle panels show the corresponding primary cell cultures 2 days after isolation from treated vesicles. Lower panels show the cultures after 14 days of drug-free regeneration. Arrows indicate newly formed metacestode vesicles. Scale bar, 1 mm.

**Figure 5.**
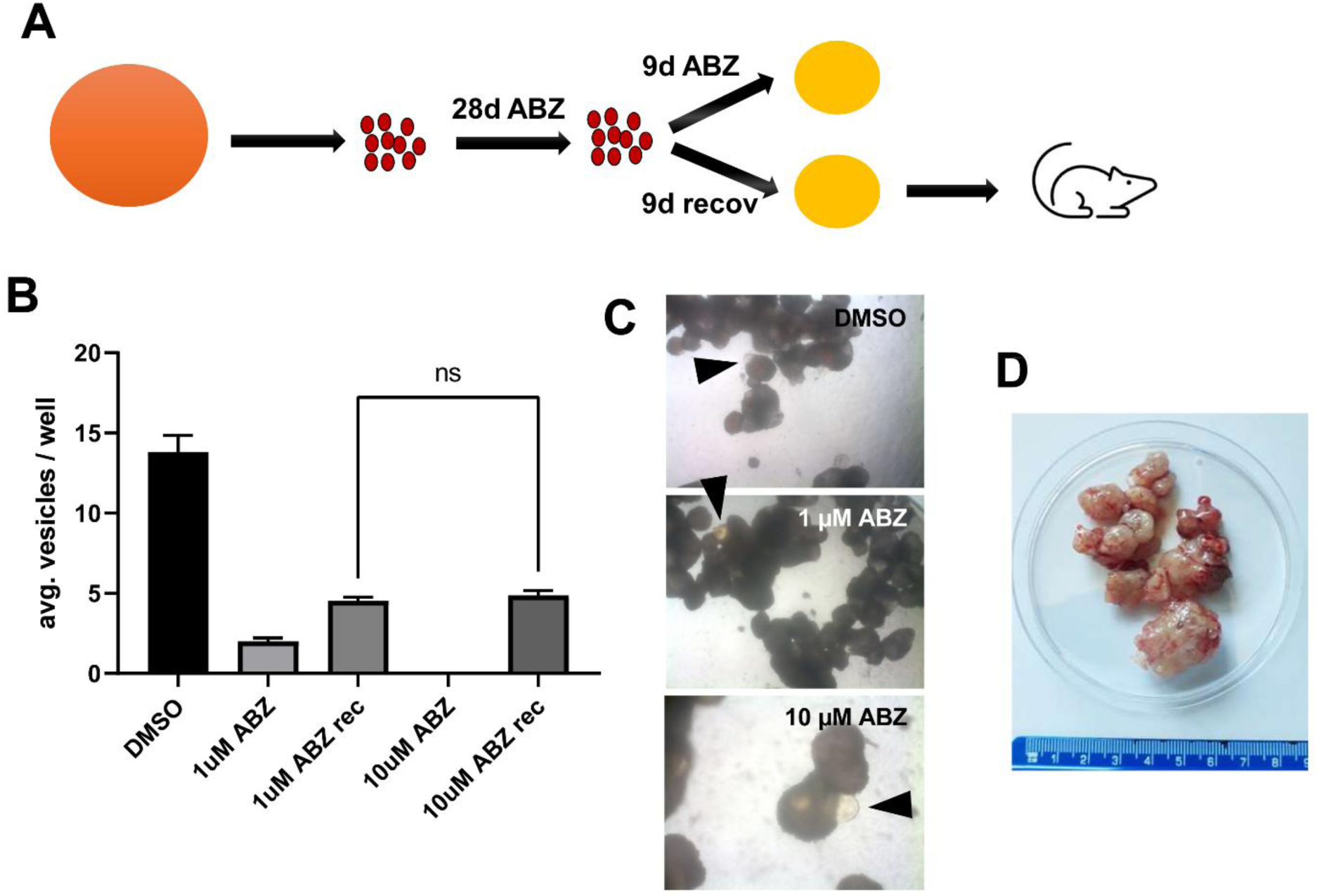
Primary cells retain regenerative capacity after prolonged ABZ exposure. (A) Experimental design. Primary cells were isolated from untreated *E. multilocularis* metacestode vesicles and cultured for 28 days in the presence of DMSO (control), 1 µM ABZ, or 10 µM ABZ. After 28 days, cultures were divided and either maintained under the respective ABZ concentration for an additional 9 days or transferred to drug-free medium for a 9-day recovery period. Vesicle formation was subsequently assessed. Primary cells recovered after treatment with 10 µM ABZ were additionally injected into a jird to assess their capacity to generate parasite tissue *in vivo*. Parasite tissue was recovered 2 months after injection. (B) Number of metacestode vesicles formed after continuous ABZ treatment or following drug withdrawal. Three independent biological replicates were analyzed. For each biological replicate, two parallel primary cell cultures were established: one was continuously maintained under the respective ABZ concentration, whereas the second was treated with ABZ for 28 days followed by 9 days of drug-free recovery. Newly formed vesicles were counted at the end of the experiment. Data are shown as mean ± SD (*n* = 3 biological replicates). The numbers of vesicles formed following drug withdrawal after previous exposure to 1 or 10 µM ABZ were compared using an unpaired two-tailed Student’s t-test; ns, not significant. (C) Representative images of primary cell cultures following the 9-day drug-free recovery period after 28 days of treatment with DMSO, 1 µM ABZ, or 10 µM ABZ. Arrowheads indicate newly formed metacestode vesicles. (D) Parasite tissue recovered from a jird 2 months after injection of primary cells that had been treated with 10 µM ABZ for 28 days followed by drug withdrawal.

**Figure 6.**
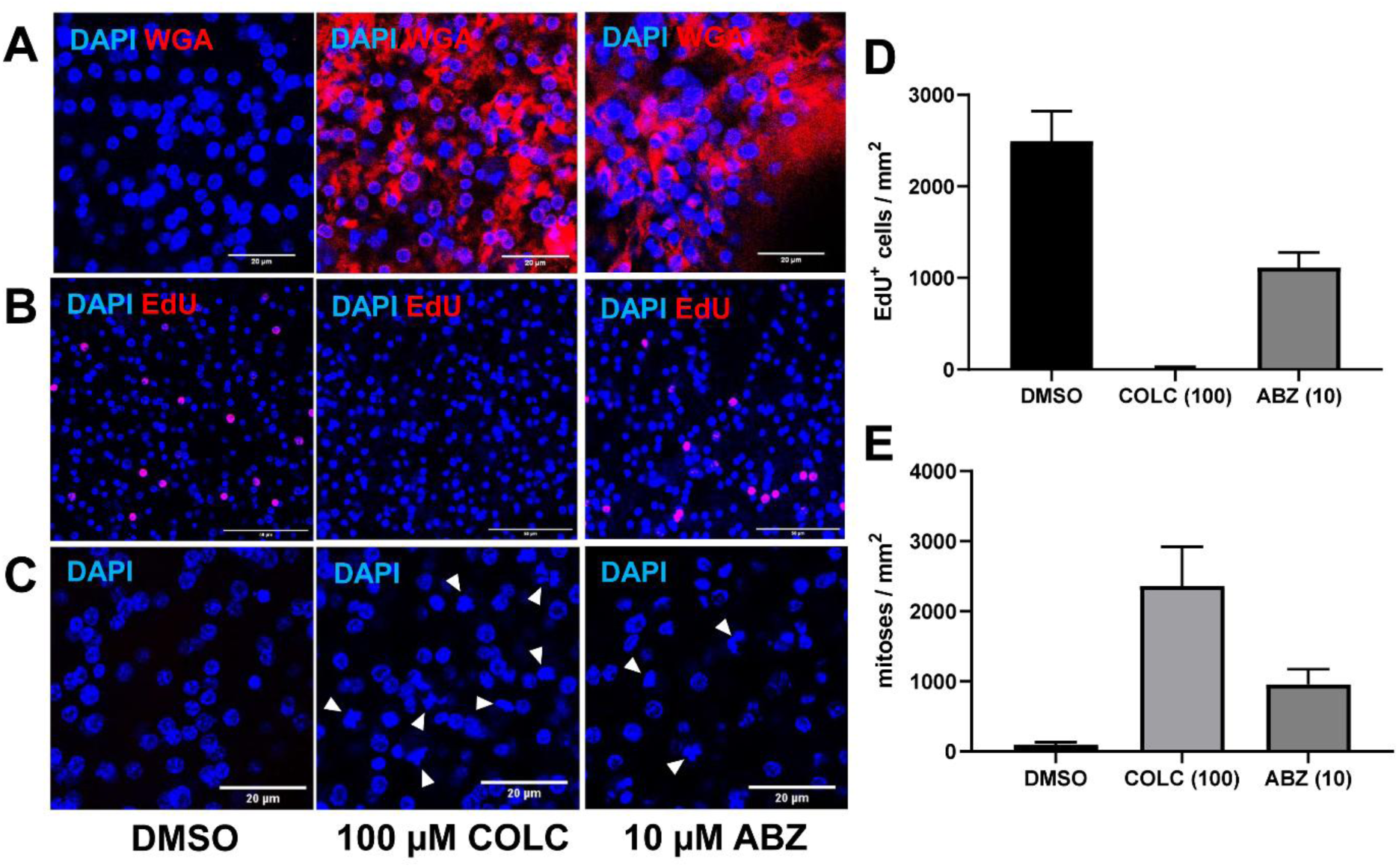
ABZ inhibits tegumental transport while allowing continued DNA synthesis and mitotic activity. Intact *E. multilocularis* metacestode vesicles were treated with DMSO, 100 µM colchicine (COLC), or 10 µM ABZ. (A) After 24 h of treatment, vesicles were stained with wheat germ agglutinin (WGA) to visualize carbohydrate-rich components transported to the parasite surface. In DMSO-treated vesicles, WGA staining was predominantly associated with the parasite surface, whereas treatment with COLC or ABZ resulted in pronounced accumulation of WGA-positive material in cytons beneath the tegument, indicative of impaired transport to the surface. Nuclei were counterstained with DAPI. (B) Representative EdU staining after 72 h of treatment. EdU-positive cells remained readily detectable following ABZ treatment, whereas they were virtually absent following COLC treatment. (C) Representative DAPI staining after 72 h of treatment showing mitotic figures (arrowheads). Mitotic cells remained detectable in ABZ-treated vesicles and accumulated strongly following COLC treatment. Mitotic figures were identified based on the characteristic morphology of condensed chromosomes in DAPI-stained nuclei. (D, E) Quantification of EdU+ cells (D) and mitotic figures (E) per mm² of germinal layer after 72 h of treatment. Five vesicles were analyzed per treatment condition, and two randomly selected microscopic fields were quantified per vesicle. Values from the two fields were averaged for each vesicle, resulting in five independent biological replicates per treatment condition. Data are shown as mean ± SD. No inferential statistical analysis was performed, as the experiment was designed to determine whether inhibition of tegumental transport by ABZ occurs under conditions in which DNA synthesis and mitotic activity remain detectable. Scale bars, 20 µm (A, C) and 50 µm (B).

**Figure 7.**
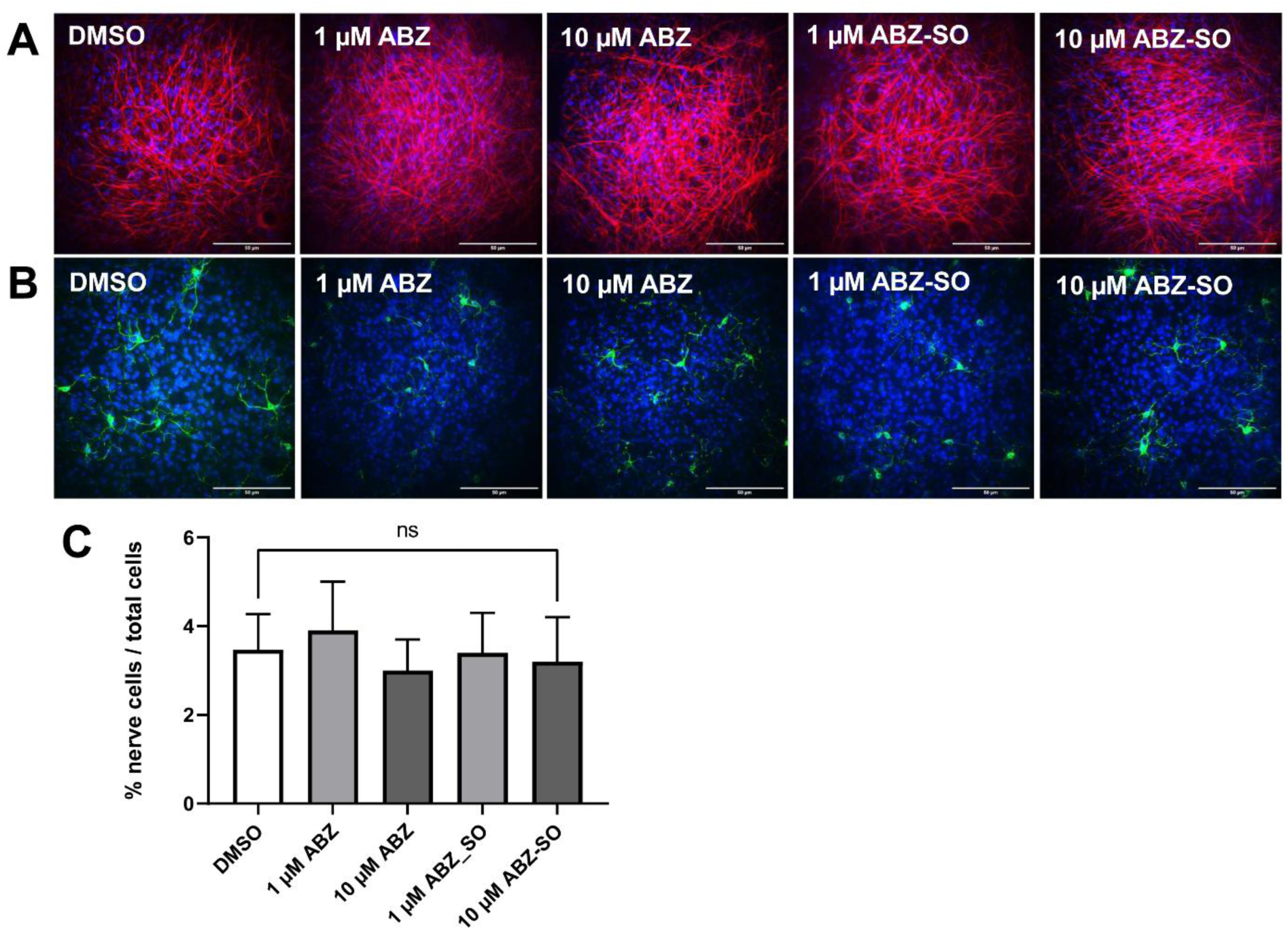
Effects of ABZ and ABZ-SO on differentiated muscle and nerve cells of *E. multilocularis* metacestode vesicles. Metacestode vesicles were treated for 10 days with DMSO (vehicle control), 1 or 10 µM ABZ, or 1 or 10 µM ABZ-SO. **(A)** Representative images of phalloidin-TRITC staining showing the organization of muscle cells and muscle fibers following the indicated treatments. **(B)** Representative images of immunofluorescence staining with an antibody against acetylated α-tubulin (acTub), visualizing nerve cells and neuronal processes under the same treatment conditions. Scale bars, 50 µm. **(C)** Quantification of nerve cells relative to the total number of cells. For each treatment condition, three independent vesicles were analyzed, with two randomly selected microscopic fields quantified per vesicle. The mean of the two fields was calculated for each vesicle, and these three vesicle means were used as individual values for statistical analysis (n = 3 vesicles per condition). Data are shown as mean ± SD.

**Figure 8.**
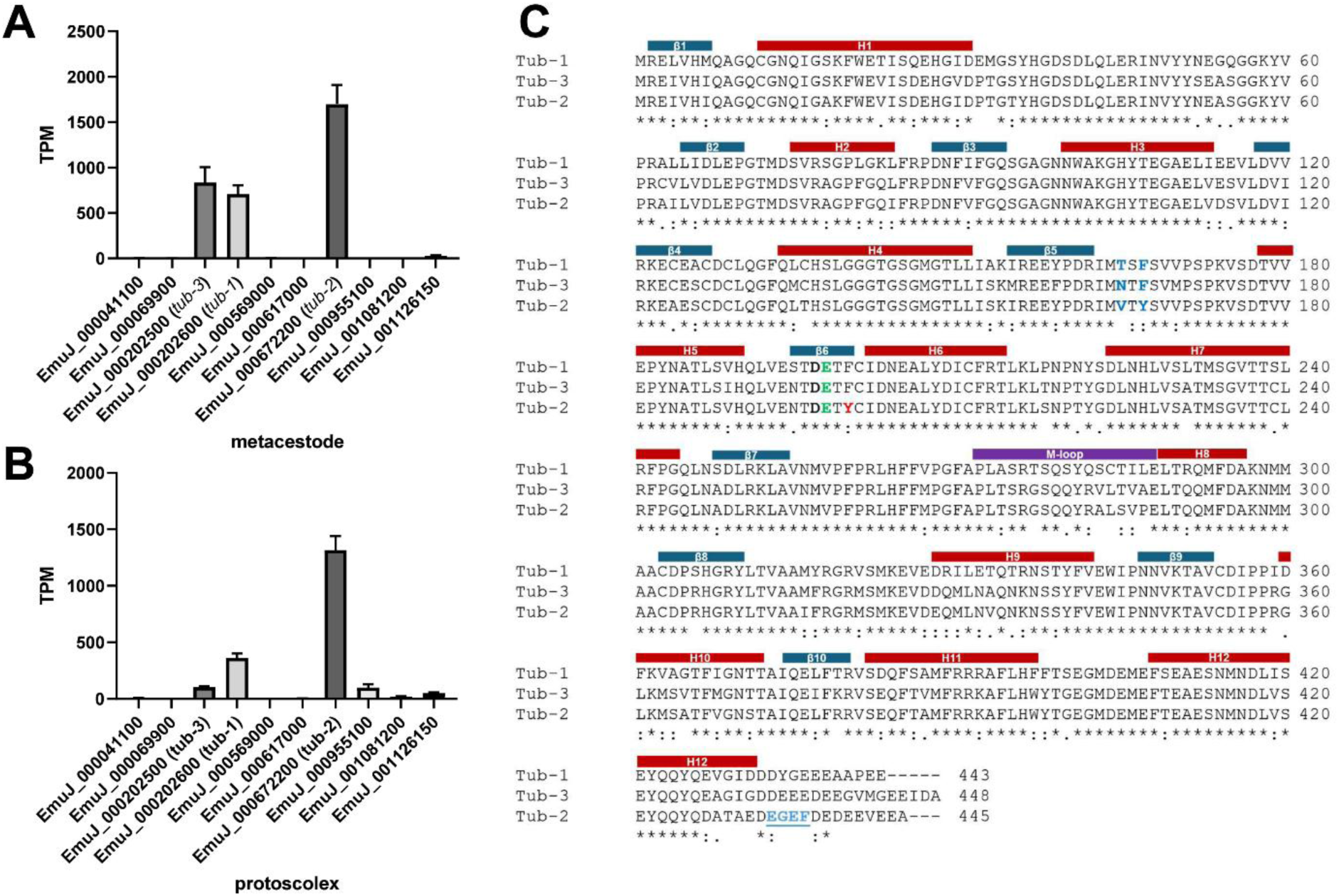
Expression and sequence characteristics of *E. multilocularis* β-tubulin isoforms. **(A, B)** Expression of the ten predicted *E. multilocularis* β-tubulin genes in metacestode vesicles (A) and protoscoleces (B). Transcript abundance is shown as transcripts per million (TPM) and is based on RNA-seq data from Herz et al. (2024). Bars represent mean TPM values ± SD from three biological replicates. The previously characterized β-tubulin isoforms *tub-1* (EmuJ_000202600), *tub-2* (EmuJ_000672200), and *tub-3* (EmuJ_000202500) are indicated. **(C)** Amino acid sequence alignment of Tub-1, Tub-2, and Tub-3 using the sequences originally described by Brehm et al. (2000). Predicted α-helices (H1–H12), β-strands (β1–β10), and the M-loop are indicated above the alignment. E198, implicated in benzimidazole binding, is highlighted in green, whereas residue 200, a major determinant of benzimidazole susceptibility, is highlighted in red; Tub-2 contains Y200, whereas Tub-1 and Tub-3 contain F200. Residues at positions 165 and 167, included in the subsequent structural analyses, are highlighted in blue. The C-terminal EGEF motif of Tub-2, corresponding to an axonemal β-tubulin epitope and recognized by the Tub2.1 antibody, is shown in blue and underlined. Sequence conservation is indicated below the alignment: asterisks indicate identical residues, colons indicate residues with strongly similar properties, and periods indicate residues with weakly similar properties.

**Figure 9.**
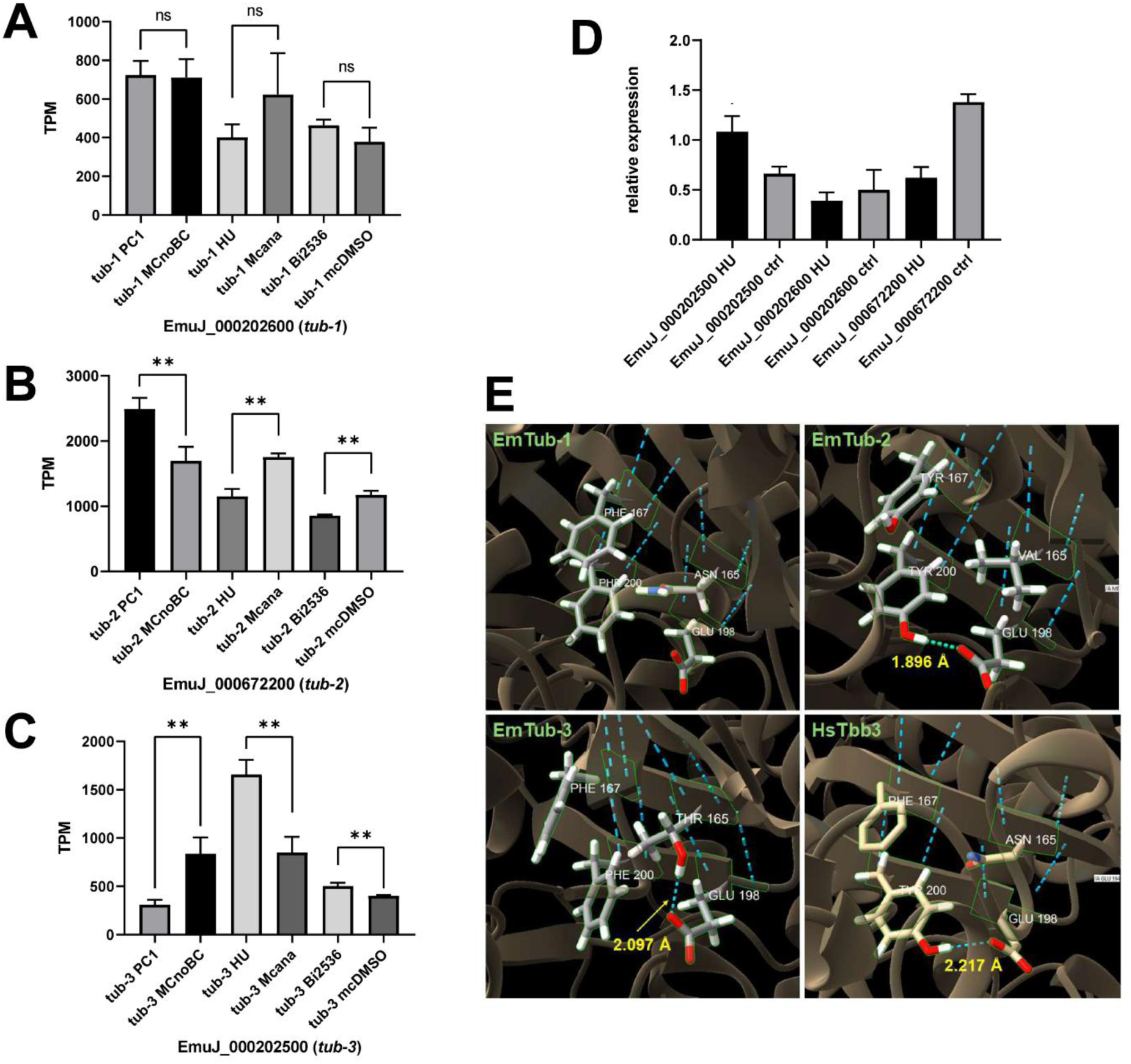
GC-associated expression of *tub-2* and structural differences among major *E. multilocularis* β-tubulin isoforms. (A–C) Relative expression of *tub-1* (EmuJ_000202600) (A), *tub-2* (EmuJ_000672200) (B), and *tub-3* (EmuJ_000202500) (C) under conditions associated with enrichment or depletion of GC. Transcript abundance is shown as transcripts per million (TPM) and was obtained from the RNA-seq datasets of Herz et al. (2024). Expression in primary cells after 2 days of cultivation (PC1), which are enriched in GC, was compared with that in metacestode vesicles without brood capsules (MCnoBC). Expression following hydroxyurea (HU) treatment, which depletes proliferating GC, was compared with the corresponding untreated metacestode control (MCana), and expression following treatment with the PLK1 inhibitor BI-2536 was compared with the corresponding vehicle-treated control (mcDMSO). Bars represent mean TPM values ± SD from three biological replicates. Individual values from the three biological replicates were used for statistical analysis using unpaired two-tailed t-tests. Statistical significance is indicated as follows: **P < 0.01; ns, not significant. (D) Independent validation of the effect of HU treatment on the expression of *tub-1*, *tub-2*, and *tub-3* by qRT-PCR. Expression in HU-treated metacestode vesicles is shown relative to the corresponding untreated controls. Bars represent mean ± SD of three technical qRT-PCR replicates from a single experiment. No inferential statistical analysis was performed because these technical replicates do not represent independent biological observations. (E) Structural environment of E198 in *E. multilocularis* Tub-1, Tub-2, and Tub-3 and human β-tubulin class III (HsTBB3). Close-up views show residues 165, 167, 198, and 200 in the predicted three-dimensional structures. Hydrogen bonds involving E198 are indicated by dashed lines, with the corresponding distances given in Å. Tub-2 contains Y200 and displays a hydrogen bond between E198 and Y200 (1.896 Å). Human TBB3, which likewise contains Y200, is shown as a reference β-tubulin and displays a corresponding E198–Y200 hydrogen bond (2.217 Å). In contrast, the F200-containing *E. multilocularis* isoforms Tub-1 and Tub-3 show distinct local interaction patterns: no corresponding E198–F200 hydrogen bond is observed in Tub-1, whereas in Tub-3 E198 forms a hydrogen bond with T165 (2.097 Å). HsTBB3 was included as a structural reference because the E198/Y200 configuration has previously been implicated in reduced benzimidazole susceptibility. Structural models were obtained from the AlphaFold Protein Structure Database and visualized and analyzed using UCSF ChimeraX.

**Figure 10.**
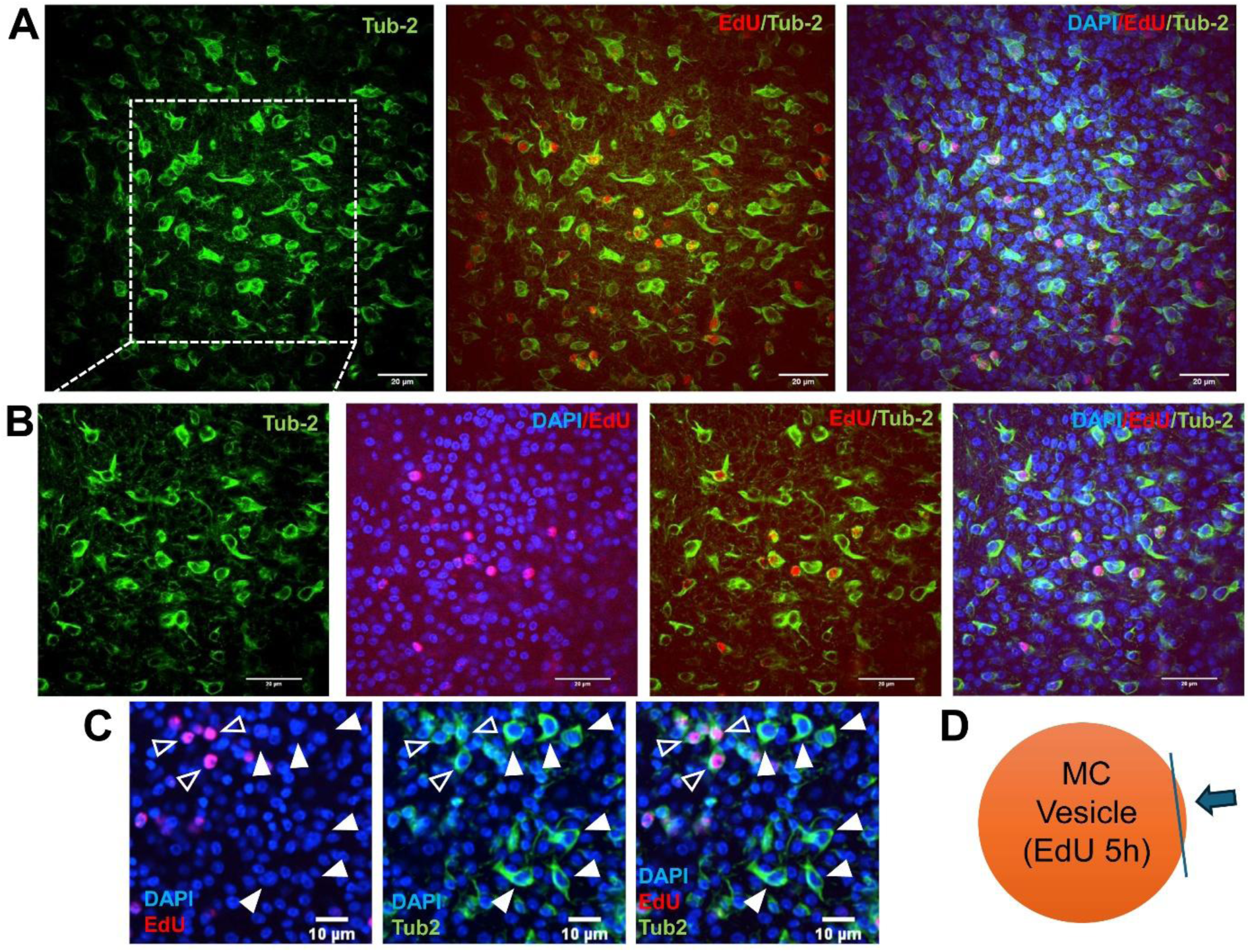
Tub-2 expression in proliferating cells of the *E. multilocularis* germinal layer. (A) Representative confocal images of the germinal layer of metacestode vesicles stained with the Tub2.1 antibody and following EdU incorporation to label proliferating cells. Three-dimensional projections are shown for Tub-2, EdU/Tub-2, and the merged channels, as indicated. (B) Higher-magnification view of the region indicated in (A), shown as single confocal optical sections. Images display Tub-2 alone, DAPI/EdU, EdU/Tub-2, and the merged channels, as indicated. DAPI was used for nuclear counterstaining. (C) Higher-magnification single confocal optical section illustrating the different Tub-2 staining patterns observed in EdU-positive and EdU-negative cells. Open arrowheads indicate EdU+/Tub-2+ cells, which generally display a less intense Tub-2 signal, whereas closed arrowheads indicate EdU-/Tub-2+ cells characterized by an intense, spindle-shaped Tub-2 staining pattern. (D) Schematic representation of an *E. multilocularis* metacestode vesicle indicating the orientation and plane of view used for imaging the germinal layer. Scale bars: 20 µm in (A) and (B); 10 µm in (C).

**Figure 11.**
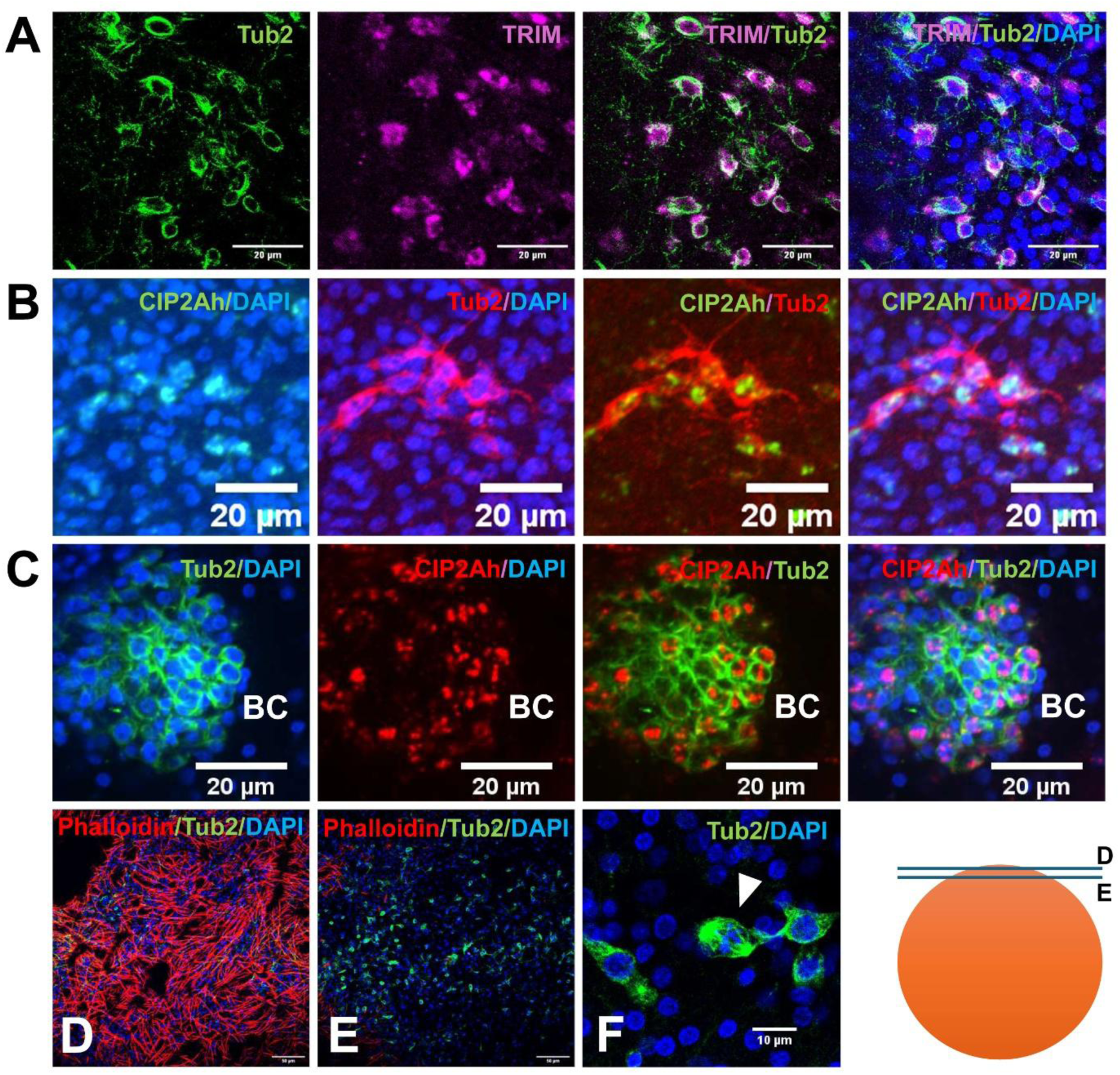
Tub-2 expression in GC of the *E. multilocularis* metacestode. (A) Colocalization of Tub-2 with the GC marker *trim* in the metacestode germinal layer. Samples were stained with the Tub2.1 antibody and analyzed by WISH for *trim* expression. Individual and merged channels are shown as indicated. *trim* has previously been established as a marker of *E. multilocularis* GC (Koziol et al., 2015). (B) Colocalization of Tub-2 with *CIP2Ah*, another marker associated with *E. multilocularis* GC (Herz et al., 2024), in the metacestode germinal layer. Individual and merged channels are shown as indicated. Note the distinct staining protocol for *CIP2Ah* (green) and Tub-2 (red). (C) Colocalization of *CIP2Ah* and Tub-2 in a developing brood capsule (BC). (D, E) Spatial distribution of Tub-2+ cells within the metacestode germinal layer. Images in (D) and (E) were obtained from the same confocal stack at different depths. The optical section in (D), immediately beneath the laminated layer, shows the superficial phalloidin-positive muscle network, whereas the section in (E), located 8 µm deeper within the germinal layer, reveals numerous Tub-2+ cells. Phalloidin and Tub-2 staining are shown together with DAPI nuclear counterstaining. The schematic on the right indicates the relative imaging planes corresponding to (D) and (E). (F) Higher-magnification single confocal optical section showing a putatively dividing Tub-2+ cell with prominent Tub-2 staining of a spindle-like structure (arrowhead), consistent with localization of Tub-2 to the mitotic spindle. DAPI was used for nuclear counterstaining. Scale bars: 20 µm in (A–C); 10 µm in (D–F).

**Figure 12.**
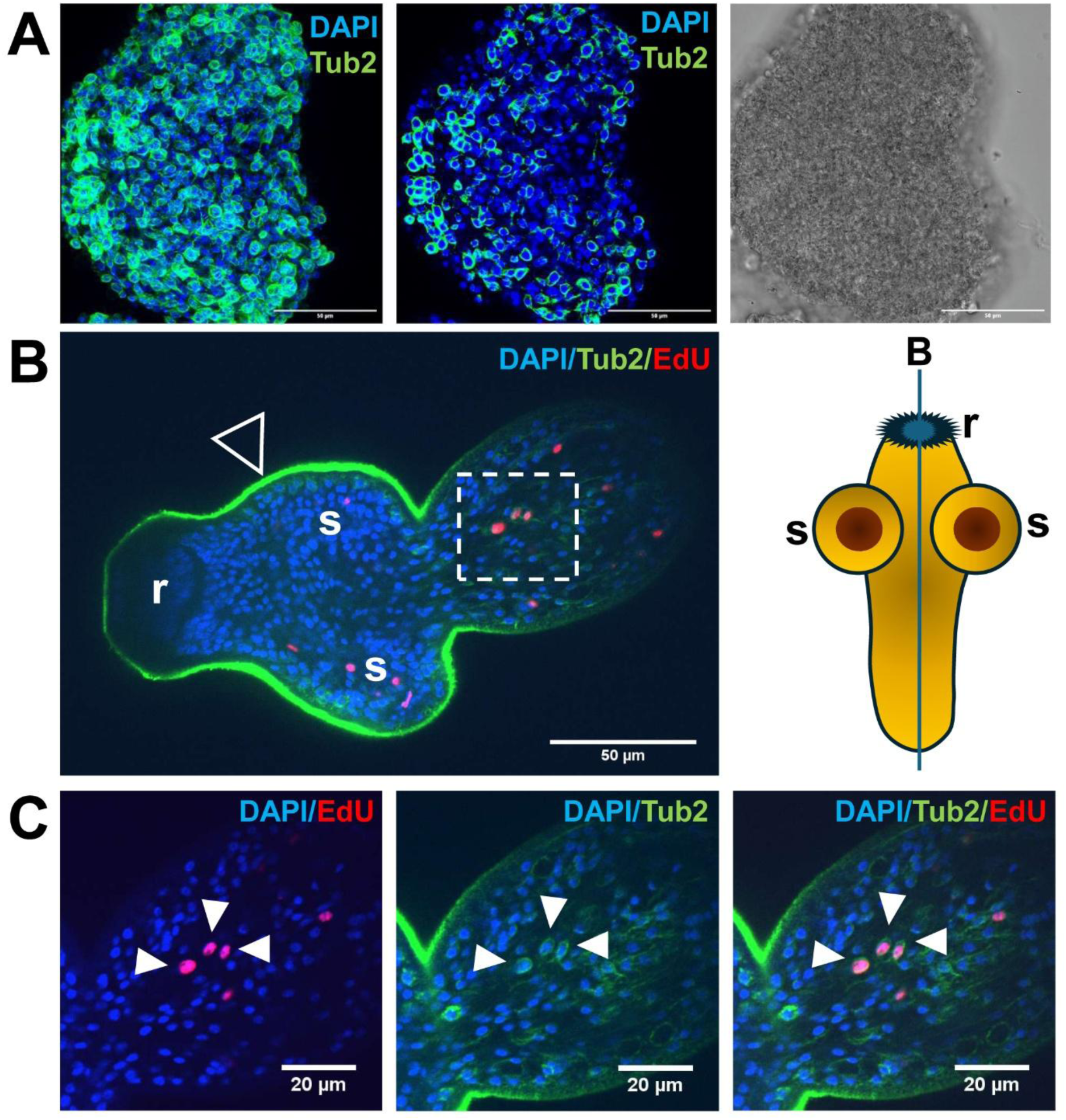
Tub-2 expression in primary cell aggregates and proliferating cells of activated protoscoleces. (A) Tub-2 expression in an aggregate derived from *E. multilocularis* primary cell culture. Tub-2 was detected by immunofluorescence (green), and nuclei were counterstained with DAPI (blue). Shown are a 3D reconstruction of the aggregate (left), a single confocal optical section (middle), and the corresponding transmitted-light image (right). Scale bars, 50 µm. (B) Tub-2 expression and proliferative activity in an activated protoscolex following a 5-h EdU pulse. Tub-2 immunofluorescence is shown in green, EdU incorporation in red, and nuclei in blue. EdU-positive cells displaying Tub-2 signal are particularly prominent in the posterior region corresponding to the prospective neck region (dashed box), which is shown at higher magnification in (C). Strong Tub-2 immunoreactivity is additionally visible along the tegument of the anterior region (open arrowhead), encompassing the suckers (s) and rostellum (r). The schematic on the right illustrates the orientation and imaging plane of the protoscolex. Scale bar, 50 µm. (C) Higher-magnification view of the posterior region indicated by the dashed box in (B), showing EdU-positive cells (red) associated with Tub-2 staining (green); nuclei are stained with DAPI (blue). Arrowheads indicate examples of EdU+ cells displaying Tub-2 signal. Scale bars, 20 µm.

**Figure 13.**
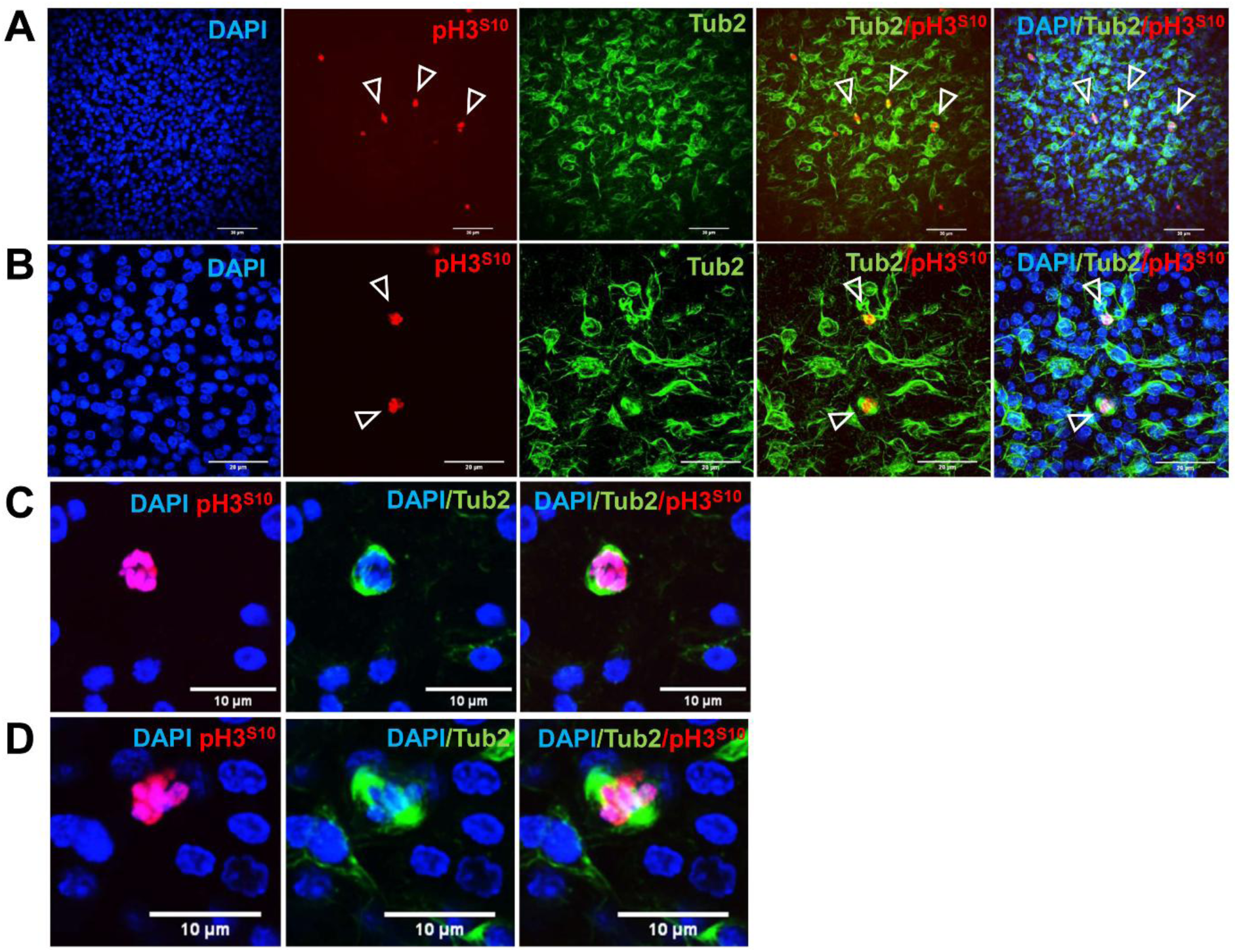
Tub-2 localizes to the mitotic spindle in *E. multilocularis* metacestode cells. Metacestode vesicles were subjected to immunohistochemical staining using the Tub-2-specific antibody Tub 2.1 (green) in combination with an antibody against histone H3 phosphorylated at serine 10 (pH3^S10^; red) as a marker of mitotic cells. Nuclei were counterstained with DAPI (blue). (A) Overview of the germinal layer shown as a 3D projection. pH3^S10^-positive mitotic cells are indicated by open arrowheads. Tub-2 staining is associated with pH3^S10^-positive cells and displays a spindle-like distribution around mitotic nuclei. Scale bars, 20 µm. (B) Higher-magnification 3D projection showing individual pH3^S10^-positive cells (open arrowheads) with prominent Tub-2 staining associated with the mitotic apparatus. Scale bars, 20 µm. (C, D) Representative single confocal slices of individual mitotic cells illustrating the characteristic spindle-like localization of Tub-2 surrounding the condensed, pH3^S10^-positive chromatin. Scale bars, 10 µm. These observations support a direct involvement of Tub-2 in mitotic spindle formation during cell division.

**Figure 14.**
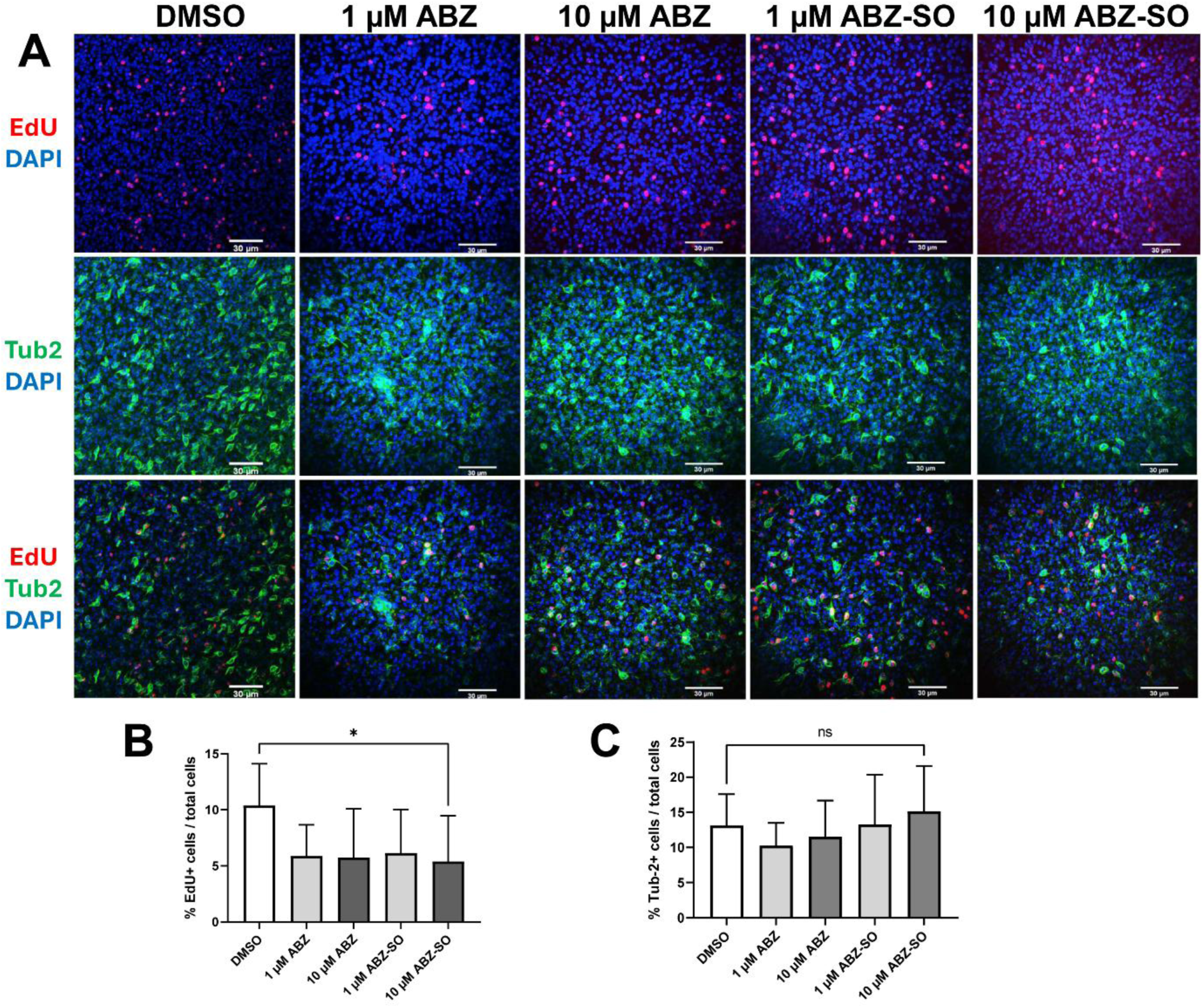
ABZ treatment reduces proliferative activity without decreasing the proportion of Tub-2+ cells in *E. multilocularis* metacestode vesicles. Intact metacestode vesicles were treated for 10 days with DMSO (control), 1 or 10 µM ABZ, or 1 or 10 µM ABZ-SO, followed by a 5-h EdU pulse. Vesicles were subsequently fixed and stained for EdU incorporation and Tub-2 using the Tub 2.1 antibody; nuclei were counterstained with DAPI. (A) Representative confocal 3D projections of vesicles following the indicated treatments. For each condition, EdU/DAPI (top), Tub-2/DAPI (middle), and merged EdU/Tub-2/DAPI (bottom) channels are shown. Scale bars, 30 µm. (B) Quantification of EdU-positive cells as a percentage of the total number of DAPI-positive cells. Treatment with either concentration of ABZ or ABZ-SO resulted in a significant reduction in the proportion of EdU-positive cells compared with the DMSO control. For clarity, a single asterisk spanning all treatment groups is shown to indicate that each individual treatment group differed significantly from the DMSO control (*P* < 0.05). (C) Quantification of Tub-2+ cells as a percentage of the total number of DAPI+ cells. In contrast to EdU incorporation, the proportion of Tub-2+ cells was not significantly affected by ABZ or ABZ-SO treatment. The single ns annotation spanning all treatment groups indicates that none of the individual treatment groups differed significantly from the DMSO control.

### Drug treatment

ABZ (Sigma-Aldrich, A4673) and ABZ sulfoxide (ABZ-SO; Cayman Chemical, 21880) were prepared as 10 mM stock solutions in DMSO. Colchicine (Sigma-Aldrich, C9754) was prepared as a 100 mM stock solution in DMSO. Imatinib was obtained from Sigma-Aldrich (CDS022173). Hydroxyurea (HU; Sigma-Aldrich, H8627) was prepared as a 2 M stock solution in DMEM without fetal calf serum. ABZ and ABZ-SO were applied at the concentrations indicated for the individual experiments. Control cultures received the corresponding concentration of DMSO. Unless stated otherwise, culture medium was exchanged every three days and freshly prepared compound was added at each medium change to maintain continuous drug exposure. For analysis of metacestode integrity, intact H95 vesicles were treated with ABZ or ABZ-SO at the indicated concentrations for seven days. Vesicle integrity was assessed morphologically, and vesicles showing loss of their characteristic intact spherical morphology were scored as damaged. Three independent biological experiments comprising three technical replicates each were performed. Long-term treatment experiments were performed with 1 or 10 µM ABZ and, where indicated, 1 or 10 µM ABZ-SO for the periods specified for the respective experiments. Colchicine (100 µM) was used at a deliberately high concentration as a positive control for pronounced microtubule disruption.

### Resazurin reduction assays

Metabolic activity of *E. multilocularis* primary cells was assessed using a resazurin reduction assay. Primary cells were seeded at 10 U per well into 96-well plates. DMEM without phenol red supplemented with 10% FCS and the respective compound was added to a final volume of 100 µl per well. Cells treated with 1% Triton X-100 served as dead controls. Plates were incubated at 37°C under a nitrogen atmosphere for the indicated treatment period. Resazurin (Sigma-Aldrich) was prepared as a 2 mg/ml stock solution in H₂O and stored at 4°C. Immediately before use, the stock solution was diluted 1:100 in PBS and 100 µl was added to each well. Fluorescence was measured immediately at 540 nm with a reference wavelength of 595 nm using a Tecan plate reader to obtain baseline values. Plates were subsequently incubated for 3 h under culture conditions before final fluorescence measurements were performed. For the experiments shown in Fig. 1, H95 primary cells were exposed to the indicated concentrations of ABZ, ABZ-SO, or imatinib for two days. Three independent biological experiments were performed, each comprising three technical replicates. Technical replicates were averaged within each experiment, and the resulting biological replicate means were used as individual values for statistical analysis.

### EdU labelling and analysis of proliferating cells

Proliferating cells were labelled with 5-ethynyl-2′-deoxyuridine (EdU) and detected according to the established protocol for *E. multilocularis* (Koziol et al., 2014; Herz et al., 2024). For EdU labeling, metacestode vesicles were incubated with 50 µM EdU for 5 h before fixation. Incorporated EdU was detected by click chemistry using a fluorescent azide according to the manufacturer’s instructions (Thermo Fisher Scientific). To determine proliferative competence following prolonged ABZ treatment, intact metacestode vesicles were continuously exposed to DMSO, 1 µM ABZ, or 10 µM ABZ for 14 days. Compounds were subsequently removed and vesicles were maintained for one additional day in drug-free medium before application of a 5-h EdU pulse. Three independent experiments were performed, using two independent preparations of isolate GH09 and one preparation of H95. Each experiment comprised three technical replicates with three vesicles per replicate. Three randomly selected microscopic fields were quantified per vesicle. Values obtained within each biological experiment were averaged, and the resulting three biological replicate means were used for statistical analysis.

For continuous-exposure experiments, primary cells isolated from untreated H95 metacestodes were maintained in DMSO, 1 µM ABZ, or 10 µM ABZ for 14 days. Cultures were subsequently either maintained for an additional seven days under the same treatment condition or transferred to drug-free medium for seven days. At the end of the resulting 21-day culture period, cells were subjected to a 5-h EdU pulse. Because prolonged primary cell cultivation resulted in heterogeneous three-dimensional aggregates, these experiments were evaluated qualitatively for the presence of EdU-positive cells rather than quantitatively.

### Regeneration assays

To determine whether GC retained developmental competence following prolonged ABZ exposure, two complementary regeneration experiments were performed using isolate H95. First, intact metacestode vesicles were treated continuously with DMSO, 1 µM ABZ, or 10 µM ABZ for 14 days. Primary cells were subsequently isolated from the treated parasite material and transferred to drug-free culture medium. Formation of new metacestode vesicles was assessed after 14 days. Three independently prepared metacestode cultures were treated and subsequently used in their entirety for primary cell isolation, yielding three independent biological replicates per condition. Second, primary cells isolated from untreated metacestodes were directly exposed to DMSO, 1 µM ABZ, or 10 µM ABZ for 28 days. Cultures were subsequently divided and either maintained for another nine days under the respective treatment condition or transferred to drug-free medium for a nine-day recovery period. Newly generated metacestode vesicles were counted at the end of the experiment. Three independent biological experiments were performed, with paired continuously treated and recovery cultures generated from each primary cell preparation.

### *In vivo* regeneration of ABZ-treated primary cells

To assess whether cells surviving prolonged ABZ exposure retained developmental competence in vivo, H95 primary cells maintained in the presence of 10 µM ABZ for 28 days were removed from drug treatment and injected intraperitoneally into a *Meriones unguiculatus*. The animal was maintained for two months before parasite tissue was recovered from the peritoneal cavity.

This study was performed in strict accordance with German (Deutsches Tierschutzgesetz, TierSchG) and European (Directive 2010/63/EU) regulations on the protection of animals. The protocol was approved by the Ethics Committee of the Government of Lower Franconia under permit number 55.2.2-2532-2-1824-10.

### Analysis of tegument-associated transport

Tegument-associated transport was analyzed using wheat germ agglutinin (WGA) according to a procedure previously established for cestode metacestodes (Guarnaschelli and Koziol, 2025). GH09 metacestode vesicles were treated for 24 h with DMSO, 10 µM ABZ, or 100 µM colchicine and subsequently fixed as described above. Following fixation, vesicles were washed with PBS containing 0.1% Triton X-100 (PBS-Tx) and permeabilized with 1% SDS for 10 min. After washing with PBS-Tx, vesicles were incubated for 4 h in PBS-Tx containing 7.5 µg/ml rhodamine-conjugated wheat germ agglutinin (WGA; RL-1022-5, Vector Laboratories, USA). Samples were subsequently washed with PBS-Tx, and nuclei were counterstained with DAPI. The distribution of WGA-positive material between the distal tegument and underlying tegumental cytons was analyzed by confocal microscopy.

To determine whether these early effects were accompanied by alterations in GC proliferation and mitosis, vesicles were treated under the same conditions for 72 h and subjected to a 5-h pulse with 50 µM EdU. Incorporated EdU was detected using the Click-iT™ EdU Alexa Fluor™ 555 Imaging Kit (Thermo Fisher Scientific) according to the manufacturer’s instructions. EdU-positive cells and mitotic figures were quantified in five vesicles per treatment condition. Two randomly selected microscopic fields were analyzed per vesicle and averaged, yielding one value per vesicle. Cell numbers were expressed per mm² of germinal layer.

### Immunohistochemistry and visualization of differentiated cell types

Whole-mount immunofluorescence of metacestode vesicles, primary cell aggregates, and activated protoscoleces was performed according to established procedures (Koziol et al., 2014; Herz et al., 2024). Tub-2 was detected using the monoclonal Anti-β-Tubulin TUBB1 antibody Tub 2.1 (Boster Biological Technology, MA1112; 1:50). Mitotic cells were detected using an antibody against histone H3 phosphorylated at serine 10 (pH3S10; Cell Signaling Technology, 9701; 1:100) according to Koziol et al. (2014). Neuronal cells and processes were visualized using the monoclonal anti-acetylated α-tubulin antibody 6-11B-1 (Santa Cruz Biotechnology, sc-23950; 1:100). Muscle fibres were visualized using TRITC-conjugated phalloidin (Sigma-Aldrich, P1951; 200 µg/ml stock solution in methanol; 1:40). For detection of Tub 2.1 and anti-acetylated α-tubulin antibodies, Alexa Fluor 488 AffiniPure Goat Anti-Mouse IgG (H+L) (Jackson ImmunoResearch, 115-545-166; 1:100) was used as secondary antibody. The pH3S10 antibody was detected using Cy3 AffiniPure Goat Anti-Rabbit IgG (H+L) (Jackson ImmunoResearch, 111-165-144; 1:100). Nuclei were counterstained with DAPI. For analysis of differentiated cells following prolonged benzimidazole exposure, GH09 metacestode vesicles were treated for ten days with DMSO, 1 or 10 µM ABZ, or 1 or 10 µM ABZ-SO and subsequently analyzed using phalloidin or anti-acetylated α-tubulin staining according to Kaethner et al. (2023b). For neuronal cell quantification, three independent vesicles were analyzed per condition and two randomly selected fields were quantified per vesicle. Values from both fields were averaged to yield one value per vesicle.

### Tub-2 immunostaining and quantitative analysis

For analysis of the relationship between Tub-2 expression and proliferative activity, H95 metacestode vesicles were subjected to a 5-h EdU (50 µM) pulse followed by fixation and Tub 2.1 immunostaining. A total of 1,432 Tub-2+ cells was analyzed to determine the proportion incorporating EdU and, conversely, the proportion of EdU+ cells displaying Tub-2 immunoreactivity. For analysis of Tub-2+ cells following benzimidazole treatment, H95 metacestode vesicles were continuously exposed for ten days to DMSO, 1 or 10 µM ABZ, or 1 or 10 µM ABZ-SO and subsequently subjected to a 5-h EdU pulse. For each condition, one randomly selected microscopic field from each of eight vesicles was quantified. The percentages of EdU+ and Tub-2+ cells relative to the total number of DAPI-positive cells were calculated.

### Whole-mount in situ hybridization and Tub-2 colocalization

Whole-mount in situ hybridization (WISH) was performed according to previously established procedures for *E. multilocularis* (Koziol et al., 2014; Herz et al., 2024). Detection of *trim* transcripts was performed using the previously described probe recognizing the repetitive *trim* sequence distributed throughout the *E. multilocularis* genome (Koziol et al., 2015). *CIP2Ah* was detected using the probe described by Herz et al. (2024). Following WISH, samples were subjected to Tub 2.1 immunostaining, and colocalization of Tub-2 with the GC markers *trim* and *CIP2Ah* was analyzed by confocal microscopy.

### Recombinant expression of *E. multilocularis* β-tubulins and validation of the Tub 2.1 antibody

The complete coding sequences of *E. multilocularis tub-1*, *tub-2*, and *tub-3* were amplified from cDNA and cloned into the pET151/D-TOPO expression vector (Invitrogen/Thermo Fisher Scientific) according to the manufacturer’s instructions. Directional cloning was achieved by inclusion of the required 5′-CACC sequence in the forward primers. The native stop codon was retained in each construct, resulting in recombinant β-tubulins carrying the vector-encoded N-terminal 6×His-V5-TEV tag while leaving the native C-termini of the proteins unchanged. Primer sequences used for amplification and cloning are listed in Table S1. The complete coding sequences of the three β-tubulin constructs and their correct reading frames were verified by DNA sequencing. Directional TOPO cloning reactions were transformed into *Escherichia coli* TOP10 cells for plasmid propagation and characterization, following the manufacturer’s protocol. Recombinant plasmids containing the complete *tub-1*, *tub-2*, or *tub-3* open reading frames in the correct orientation and reading frame were verified by DNA sequencing. Sequence-verified expression constructs were subsequently introduced into *E. coli* BL21 Star(DE3) for recombinant protein expression. Protein expression was induced with IPTG according to the manufacturer’s protocol for the Champion pET151 Directional TOPO Expression System.

His-tagged recombinant β-tubulins were purified from bacterial lysates by affinity chromatography, separated by SDS-PAGE, and analyzed by Western blotting. The identity of the purified recombinant proteins was confirmed using an anti-V5 antibody (Invitrogen, 460705; 1:5,000), whereas the isoform specificity of Tub 2.1 was assessed using the monoclonal anti-β-tubulin antibody Tub 2.1 (Boster Biological Technology, MA1112; 1:50). Detection was performed using an HRP-conjugated anti-mouse secondary antibody (Dianova, 115-035-044; 1:10,000) according to the Western blotting procedure described previously (Kaethner et al., 2023b).

### RNA isolation and quantitative RT-PCR

HU treatment, RNA isolation, cDNA synthesis, and quantitative real-time PCR were performed according to the procedures described previously (Herz et al., 2024). The same HU treatment regime used in that study was employed to deplete proliferating GC. Relative transcript levels of *tub-1* (EmuJ_000202600), *tub-2* (EmuJ_000672200), and *tub-3* (EmuJ_000202500) were determined using the primers listed in Table S1. Expression was normalized against *elp*, as established previously for *E. multilocularis* (Herz et al., 2024). Quantitative PCR was performed using a StepOnePlus Real-Time PCR System (Applied Biosystems). Each reaction contained 2.4 µl 5× HOT FIREPol EvaGreen qPCR Mix Plus (1× final concentration), 0.72 µl of each primer (300 nM final concentration), 6.96 µl DNase-free water, and cDNA. Cycling conditions consisted of an initial incubation for 15 min at 95°C followed by 40 cycles of 15 s at 95°C, 20 s at 58 or 60°C, depending on the primer pair, and 20 s at 72°C. Fluorescence data were acquired at 72°C. The experiment shown in Fig. 9D was performed once with three technical qRT-PCR replicates. Accordingly, these measurements were used to assess the direction and magnitude of expression changes but were not treated as independent biological replicates for inferential statistical analysis.

### Analysis of previously published transcriptome data

β-Tubulin expression was analyzed using the previously published RNA-seq datasets of Herz et al. (2024). Transcript abundance was expressed as transcripts per million (TPM). Expression of the ten predicted *E. multilocularis* β-tubulin genes was compared between metacestode vesicles and protoscoleces. Expression of *tub-1*, *tub-2*, and *tub-3* was additionally analyzed in GC-enriched primary cultures, HU-treated metacestodes, and metacestodes treated with the PLK1 inhibitor BI-2536 together with their corresponding control conditions. Three biological replicates from the original datasets were analyzed for each condition.

### β-Tubulin sequence and structural analyses

Protein sequences of *E. multilocularis* Tub-1, Tub-2, and Tub-3 were retrieved from UniProt using the accession numbers corresponding to the β-tubulin sequences originally described by Brehm et al. (2000). Multiple sequence alignment was performed using Clustal Omega via the EMBL-EBI Job Dispatcher. Residues 165, 167, 198, and 200 were specifically examined because of their established association with benzimidazole susceptibility. Assignment of α-helices, β-strands, and the M-loop was based on the AlphaFold structural information provided through UniProt. Predicted three-dimensional structures of *E. multilocularis* Tub-1, Tub-2, and Tub-3 and human β-tubulin class III (HsTBB3) were obtained from the AlphaFold Protein Structure Database and visualized and analyzed using UCSF ChimeraX version 1.10.1. The local structural environment surrounding residues 165, 167, 198, and 200 was examined. Hydrogen bonds involving E198 and neighboring residues were identified using the hydrogen-bond detection algorithm implemented in ChimeraX, and distances between interacting atoms were determined directly from the predicted structures.

### Confocal microscopy and image analysis

Fluorescence samples were analyzed using a Leica TCS SP5 confocal microscope (Leica Microsystems, Wetzlar, Germany) and/or a Nikon Eclipse Ti2-E confocal microscope (Nikon, Düsseldorf, Germany), as appropriate for the respective experiments. Image acquisition parameters were kept constant between treatment groups within experiments involving quantitative comparisons. Three-dimensional projections and single optical sections were generated and images were analyzed using Fiji/ImageJ. Cell counting and measurement of analyzed areas were likewise performed using Fiji/ImageJ. For quantitative analyses, microscopic fields were randomly selected within the germinal layer. DAPI-positive nuclei were used to determine total cell numbers where indicated.

### Statistical analysis

Statistical analyses were performed using GraphPad Prism version 9 (GraphPad Software, San Diego, CA, USA). Data are presented as mean ± standard deviation (SD). Biological replicates or individual vesicles, depending on the experimental design, were used as independent observations as specified in the corresponding figure legends. Technical replicates were averaged before statistical analysis and were not treated as independent biological observations. Comparisons between treatment groups and their respective controls were performed using unpaired two-tailed Student’s *t*-tests as indicated in the figure legends. A *P* value < 0.05 was considered statistically significant. No inferential statistical test was applied to experiments for which only technical rather than independent biological replicates were available.

## Results

### ABZ affects metacestode vesicles but not the metabolic activity of GC-enriched primary cultures

We first assessed the effects of ABZ and its major active metabolite ABZ-SO on intact *E. multilocularis* metacestode vesicles. Previous *in vitro* studies had demonstrated pronounced effects of benzimidazoles on metacestode vesicle integrity under different experimental exposure regimens, including prolonged incubation with relatively high initial drug concentrations (Jura et al., 1998; Ingold et al., 1999; Küster et al., 2014). Since the subsequent experiments in this study were designed to assess parasite responses under sustained drug exposure, we first established the effects of ABZ and ABZ-SO on intact vesicles under the same treatment conditions, employing an *in vitro* culture system for metacestode vesicles previously established by us (Spiliotis et al., 2008; Spiliotis and Brehm, 2009). Fresh compound was added at each medium change every three days throughout the treatment period. Following seven days of treatment, both compounds induced pronounced concentration-dependent damage to the vesicles, as determined by loss of vesicle integrity (Fig. 1A, B). These results confirmed the expected anti-parasitic activity of ABZ and ABZ-SO against the metacestode under the sustained exposure conditions employed throughout this study.

We next asked whether ABZ and ABZ-SO also affected parasite primary cells. Freshly isolated *E. multilocularis* primary cell cultures are strongly enriched (>80%) in GC, the only proliferative somatic cell population of the metacestode (Koziol et al., 2014; Herz et al., 2024), and have previously provided a useful experimental system to investigate drug effects on the parasite stem cell compartment (Schubert et al., 2014; Koike et al., 2022). Primary cells were therefore exposed to ABZ or ABZ-SO immediately after isolation, and metabolic activity was determined by resazurin reduction after two days of culture (Fig. 1D). Metabolic viability assays are widely used to assess cellular responses to benzimidazoles and other microtubule-targeting compounds, including their effects on mammalian cancer cells (Pourgholami et al., 2005; Ghasemi et al., 2017; Jung et al., 2022). As a positive control, we used imatinib, which we previously demonstrated to impair *E. multilocularis* primary cells and metacestode development (Hemer and Brehm, 2012). Under the present assay conditions, imatinib caused a pronounced reduction in resazurin conversion (Fig. 1C), demonstrating that drug-induced impairment of primary cell metabolic activity could readily be detected.

In contrast, neither ABZ nor ABZ-SO significantly affected resazurin conversion after two days of treatment over the concentration range tested (Fig. 1D). ABZ activity against *E. multilocularis* primary cells has previously been reported using a longer, five-day exposure period (Kaethner et al., 2023a). Although the different treatment periods preclude a direct comparison between the effects observed in intact vesicles and primary cell cultures, the absence of a detectable short-term effect on the GC-enriched primary cultures contrasted with the pronounced damage observed in intact metacestode vesicles. This observation prompted us to investigate more directly how ABZ treatment affects the proliferative GC compartment of the metacestode.

### GC retain proliferative capacity following prolonged ABZ treatment

The lack of a detectable effect of ABZ and ABZ-SO on the metabolic activity of GC-enriched primary cultures prompted us to directly investigate the proliferative capacity of GC following prolonged drug exposure. Incorporation of the thymidine analogue 5-ethynyl-2′-deoxyuridine (EdU) provides a direct measure of DNA synthesis and is routinely used to identify proliferating GC in *E. multilocularis*. In previous studies, we and others have employed EdU incorporation to quantify changes in GC proliferation following pharmacological interference with parasite signalling and cell-cycle pathways (Herz et al., 2024; Kaethner et al., 2023a; Koike et al., 2022; Koziol et al., 2014; Tian et al., 2024; Wang et al., 2026). We therefore used short-term EdU incorporation to determine whether GC retained their capacity to proliferate after prolonged ABZ exposure.

Intact metacestode vesicles were treated with 1 or 10 µM ABZ for 14 days. Following removal of the drug and a one-day recovery period in drug-free medium, the vesicles were subjected to a 5-h EdU pulse, and the proportion of EdU+ nuclei relative to the total number of DAPI+ nuclei was determined (Fig. 2A,B,C). Treatment with 1 µM ABZ, a concentration in the range of plasma concentrations reported for the active metabolite ABZ-SO in treated patients (Lötsch et al., 2016), did not significantly alter the proportion of EdU+ cells compared with the DMSO control (Fig. 2B). In contrast, treatment with 10 µM ABZ, a concentration substantially exceeding typical clinically achieved ABZ-SO levels, resulted in a significant reduction in the proportion of EdU+ cells. Importantly, however, EdU+ cells remained readily detectable even after 14 days of treatment with 10 µM ABZ (Fig. 2B, C). At this time point, the treated vesicles had lost their structural integrity and displayed a collapsed morphology that would commonly be interpreted as parasite death in morphology-based viability assays. Thus, even severely damaged metacestode tissue retained a population of germinative cells capable of re-entering proliferation after drug withdrawal.

Thus, prolonged exposure to a high concentration of ABZ reduced the subsequent proliferative activity of the GC population but did not abolish it. Since EdU incorporation was assessed after a one-day drug-free recovery period, these experiments do not establish whether the remaining GC were capable of proliferating in the continuous presence of ABZ. Rather, they demonstrate that proliferatively competent GC persisted after prolonged ABZ exposure and resumed or maintained DNA synthesis following drug removal. Together with the lack of a detectable short-term effect of ABZ on GC-enriched primary cultures, these observations prompted us to investigate whether primary cells retain their proliferative capacity during prolonged and continuous exposure to ABZ.

### Proliferative cells persist during continuous long-term ABZ exposure

To determine whether proliferative cells persist not only after ABZ withdrawal but also during continuous drug exposure, we next employed GC-enriched primary cultures. Primary cells isolated from untreated metacestode vesicles were cultured in the presence of DMSO, 1 µM ABZ, or 10 µM ABZ for 14 days. Cultures were then either maintained for an additional 7 days in the presence of the respective ABZ concentration or transferred to drug-free medium for a 7-day recovery period. At the end of the resulting 21-day culture period, proliferative activity was assessed by a 5 h EdU pulse (Fig. 3A).

EdU+ cells were readily detectable in primary cell aggregates following the 7-day recovery period after treatment with both 1 and 10 µM ABZ (Fig. 3B). Importantly, EdU+ cells were also present in cultures that had been continuously exposed to ABZ for the entire 21-day period, including cultures maintained at 10 µM ABZ. Thus, DNA-synthesizing cells persisted even during prolonged and uninterrupted exposure to a high ABZ concentration.

Because primary cells progressively form heterogeneous three-dimensional aggregates during prolonged culture, reliable quantitative determination of the proportion of EdU+ cells was not feasible in this experimental setting. We therefore restricted this experiment to a qualitative assessment of the presence or absence of proliferatively active cells. Nevertheless, the persistence of EdU incorporation during continuous ABZ exposure, together with the recovery experiments and the quantitative analysis of intact metacestodes described above, provides further evidence that proliferative GC are not eliminated by prolonged ABZ treatment.

### ABZ treatment does not impair the regenerative capacity of GC

The persistence of EdU+ cells during prolonged ABZ exposure indicated that proliferative GC were not eliminated by the drug. However, EdU incorporation alone does not establish whether these cells retain their developmental potential. We therefore investigated whether GC persisting after prolonged ABZ treatment remained capable of regenerating metacestode tissue.

Intact metacestode vesicles were treated with DMSO, 1 µM ABZ, or 10 µM ABZ for 14 days. Primary cells were subsequently isolated from the treated vesicles and transferred to drug-free medium, in which their capacity to regenerate metacestode vesicles was assessed over an additional 14-day period (Fig. 4A). Despite the pronounced damage induced by ABZ in intact metacestodes (>80% lost structural integrity during these 14 days), primary cells could be recovered from vesicles exposed to both ABZ concentrations and established primary cell cultures (Fig. 4C).

Most importantly, primary cells isolated from ABZ-treated metacestode vesicles retained their capacity to generate new vesicles. After 14 days of drug-free culture, newly formed vesicles were observed in cultures derived from DMSO-treated controls as well as from metacestodes previously exposed to 1 or 10 µM ABZ (Fig. 4C). Quantification revealed no significant difference in the number of newly generated vesicles between either ABZ treatment group and the DMSO control (Fig. 4B).

Thus, even prolonged exposure of intact metacestodes to 10 µM ABZ did not detectably impair the subsequent regenerative capacity of the recovered primary cells. Together with the persistence of EdU+ cells during continuous ABZ exposure, these findings demonstrate that ABZ treatment does not eliminate the proliferative and developmentally competent GC population. These findings prompted us to determine whether the regenerative capacity of GC is maintained even after substantially longer periods of direct ABZ exposure.

### GC retain regenerative capacity after extended direct ABZ exposure

The regenerative capacity of primary cells isolated from ABZ-treated metacestodes suggested that developmentally competent GC persist after prolonged drug exposure. We therefore asked whether this capacity is also retained when GC-enriched primary cultures are subjected directly to substantially longer periods of ABZ treatment.

Primary cells isolated from untreated metacestode vesicles were cultured in the continuous presence of DMSO, 1 µM ABZ, or 10 µM ABZ for 28 days. Cultures were then divided and either maintained for an additional 9 days under the respective drug condition or transferred to drug-free medium for a 9-day recovery period (Fig. 5A). Regenerative capacity was assessed by monitoring the formation of new metacestode vesicles.

Continuous exposure to ABZ markedly reduced vesicle formation, and no newly formed vesicles were detected in cultures continuously maintained at 10 µM ABZ (Fig. 5B). Importantly, however, withdrawal of ABZ after 28 days of treatment resulted in renewed vesicle formation. Following the 9-day drug-free recovery period, primary cells previously exposed to either 1 or 10 µM ABZ generated new metacestode vesicles (Fig. 5B, C). The numbers of vesicles formed after drug withdrawal did not differ significantly between cultures previously exposed to 1 or 10 µM ABZ, demonstrating that cells with regenerative capacity persisted even after 28 days of continuous exposure to 10 µM ABZ.

To further test whether these cells retained developmental competence *in vivo*, primary cells exposed to 10 µM ABZ for 28 days were withdrawn from drug treatment for 9 days and injected into a jird. Two months after injection, parasite tissue could be recovered from the animal (Fig. 5D), demonstrating that cells surviving prolonged high-dose ABZ exposure retained the capacity to regenerate parasite tissue *in vivo*.

Together, these experiments show that extended ABZ exposure can strongly suppress metacestode formation but fails to eradicate the developmentally competent cell population. Upon drug withdrawal, regenerative growth resumes even after 28 days of exposure to 10 µM ABZ. These findings further support the presence of an ABZ-refractory GC population and provide a potential cellular explanation for parasite regrowth following cessation of benzimidazole treatment.

### ABZ rapidly impairs tegumental transport while GC remain proliferatively active

Our finding that prolonged ABZ exposure failed to eliminate the regenerative GC population raised the question of which cellular compartments account for the pronounced effects of ABZ on intact metacestode vesicles. Recent work in the model cestode *Mesocestoides corti* identified the tegument as a particularly sensitive cellular target of benzimidazoles. Guarnaschelli and Koziol (2025) demonstrated that low concentrations of ABZ and ABZ-SO disrupt the highly organized microtubule network of the distal tegument and cause an accumulation of secretory material within the underlying tegumental cytons, consistent with impaired microtubule-dependent transport to the parasite surface. We therefore investigated whether ABZ similarly interferes with tegumental transport in *E. multilocularis* and, importantly, whether such effects occur under conditions in which GC remain proliferatively active.

Intact metacestode vesicles were treated with DMSO, 10 µM ABZ, or 100 µM colchicine. Because colchicine displays comparatively low potency against cestode microtubules (Guarnaschelli and Koziol, 2025), it was used at the deliberately high concentration of 100 µM as a positive control for pronounced microtubule disruption. After 24 h, vesicles were stained with wheat germ agglutinin (WGA) to visualize carbohydrate-rich material transported to the parasite surface (Fig. 6A). In DMSO-treated vesicles, WGA staining was predominantly associated with the parasite surface. In contrast, treatment with either ABZ or colchicine resulted in pronounced accumulation of WGA-positive material within cytons underlying the tegument, consistent with impaired transport to the distal tegument.

We next determined whether this early tegumental effect occurred while GC remained proliferatively active. After 72 h of treatment, EdU+ cells were still readily detectable in ABZ-treated vesicles, although their number was reduced compared with DMSO-treated controls (Fig. 6B, D). In contrast, EdU+ cells were virtually absent following colchicine treatment. Mitotic figures also remained readily detectable after ABZ treatment (Fig. 6C, E), whereas colchicine induced a pronounced accumulation of mitotic cells, consistent with mitotic arrest following microtubule disruption.

Thus, in agreement with the findings in *M. corti*, ABZ rapidly interfered with tegumental transport in *E. multilocularis*. Importantly, this effect occurred under conditions in which GC continued DNA synthesis and mitotic activity. These findings indicate a marked difference in the susceptibility of tegument-associated microtubule-dependent processes and the proliferative GC compartment to ABZ.

### Differentiated muscle and nerve cells persist during prolonged ABZ treatment

Having observed an early effect of ABZ on tegumental transport, we next investigated whether prolonged benzimidazole treatment similarly affected other differentiated cell populations of the metacestode. Intact vesicles were treated for 10 days with DMSO, 1 or 10 µM ABZ, or 1 or 10 µM ABZ-SO, followed by visualization of muscle cells using phalloidin-TRITC and nerve cells using antibodies against acetylated α-tubulin (Fig. 7A, B).

Despite prolonged exposure to ABZ or ABZ-SO, the extensive network of muscle fibers remained readily detectable under all treatment conditions, with no obvious loss of overall muscle organization (Fig. 7A). Similarly, acetylated α-tubulin-positive nerve cells and neuronal processes remained detectable following treatment with either compound (Fig. 7B). Quantification of nerve cells relative to the total cell population revealed no significant differences between the treatment groups and the DMSO control (Fig. 7C).

These findings indicate that the pronounced effects of ABZ on metacestode integrity are not associated with a general depletion of differentiated muscle or nerve cells. Together with the rapid impairment of tegumental transport observed in Fig. 6, the data instead point towards differential susceptibility of individual cellular compartments and microtubule-dependent processes within the metacestode.

### The metacestode expresses distinct β-tubulin isoforms with different benzimidazole susceptibility-associated residues

The differential effects of ABZ on microtubule-dependent processes in the metacestode prompted us to investigate the β-tubulin repertoire of *E. multilocularis*. Three genes encoding β-tubulin isoforms, *tub-1*, *tub-2*, and *tub-3*, were originally cloned and characterized from *E. multilocularis* more than two decades ago (Brehm et al., 2000). The availability of the complete parasite genome (Tsai et al., 2013) and stage-specific transcriptome data (Herz et al., 2024) now allowed us to reassess these isoforms in the context of the entire *E. multilocularis* β-tubulin gene family.

The *E. multilocularis* genome contains ten predicted β-tubulin genes (Tsai et al., 2013). Analysis of our previously published RNA-seq data (Herz et al., 2024) revealed a strongly stage-dependent expression pattern (Fig. 8A, B). In metacestode vesicles, expression was dominated by the three previously characterized isoforms *tub-1* (EmuJ_000202600), *tub-2* (EmuJ_000672200), and *tub-3* (EmuJ_000202500), whereas the remaining β-tubulin genes showed little or no detectable expression (Fig. 8A). Among these, *tub-2* was by far the most abundantly expressed β-tubulin gene. Protoscoleces showed a broader β-tubulin expression profile, although *tub-2* also represented the predominant transcript in this stage (Fig. 8B).

We next compared the amino acid sequences of Tub-1, Tub-2, and Tub-3, focusing on residues associated with benzimidazole binding and susceptibility (Fig. 8C). All three isoforms contain E198, a residue implicated in direct interactions with benzimidazole compounds (Aguayo-Ortiz et al., 2013a, b). In contrast, they differ at residue 200, one of the best-characterized determinants of benzimidazole susceptibility in β-tubulins (von Samson-Himmelstjerna et al., 2007; Aguayo-Ortiz et al., 2013b). Tub-1 and Tub-3 contain phenylalanine at this position (F200), whereas Tub-2 contains tyrosine (Y200). Additional sequence differences are present at residues 165 and 167. Residue 167 represents another established site associated with benzimidazole resistance, whereas residue 165 has been implicated in interactions within the benzimidazole-binding region (von Samson-Himmelstjerna et al., 2007; Aguayo-Ortiz et al., 2013a, b). These residues were therefore included in our subsequent structural analyses.

Tub-2 is further distinguished by a C-terminal EGEF motif corresponding to an axonemal β-tubulin epitope previously identified in the original characterization of the *E. multilocularis* β-tubulins (Brehm et al., 2000). This sequence provides an isoform-specific epitope recognized by the Tub2.1 antibody and thus enabled us to investigate the cellular distribution of Tub-2 within the parasite (see below).

Taken together, these data identified Tub-2 as encoded by the predominant β-tubulin transcript of the metacestode and revealed sequence characteristics that distinguish it from the co-expressed Tub-1 and Tub-3 isoforms. We therefore next investigated which cells of the metacestode express Tub-2 and whether its cellular distribution could provide a link between β-tubulin isoform usage and the differential susceptibility of parasite cell populations to ABZ.

### Tub-2 expression is associated with the proliferative GC compartment and displays a resistance-associated structural configuration

The predominance of Tub-2 in the metacestode was particularly intriguing in light of the presence of Y200 in this isoform. Amino acid substitutions at positions 167, 198, and 200 of β-tubulin are well-established determinants of benzimidazole susceptibility, with F200Y representing one of the most frequently described resistance-associated substitutions (von Samson-Himmelstjerna et al., 2007; Dilks et al., 2020). Structural modelling and molecular dynamics studies have further suggested that these residues form part of a common interaction network within the benzimidazole-binding region (Aguayo-Ortiz et al., 2013a). In particular, E198 has been identified as an important residue directly interacting with benzimidazoles, whereas introduction of Y200 has been proposed to permit formation of an intramolecular hydrogen bond between Y200 and E198. This interaction may alter the local geometry and electrostatic environment of E198 and thereby contribute to reduced benzimidazole susceptibility (Aguayo-Ortiz et al., 2013b). We therefore investigated whether the Y200-containing *E. multilocularis* Tub-2 isoform is associated with the GC compartment and whether it displays a corresponding local structural configuration.

We first re-analyzed our previously published transcriptome datasets (Herz et al., 2024), comparing expression of *tub-1*, *tub-2*, and *tub-3* under conditions associated with enrichment or depletion of proliferating GC. Expression of *tub-1* did not differ significantly between GC-enriched primary cultures and metacestode vesicles, between hydroxyurea (HU)-treated and untreated metacestodes, or following treatment with the PLK1 inhibitor BI-2536 (Fig. 9A). In contrast, *tub-2* displayed a consistent association with the proliferative GC compartment. Its expression was significantly higher in GC-enriched primary cultures than in metacestode vesicles and was significantly reduced following HU treatment, which depletes proliferating GC, as well as following BI-2536 treatment, which interferes with GC proliferation (Fig. 9B). *tub-3* showed a largely reciprocal expression pattern, with significantly lower expression in GC-enriched primary cultures and increased expression following HU treatment (Fig. 9C).

To independently validate the association between β-tubulin isoform expression and depletion of proliferating GC, we analyzed *tub-1*, *tub-2*, and *tub-3* expression by qRT-PCR following HU treatment. Consistent with the transcriptome data, qRT-PCR analysis showed reduced *tub-2* expression following HU treatment, whereas *tub-3* expression increased and *tub-1* expression remained largely unchanged (Fig. 9D). Together, these independent expression analyses strongly associated *tub-2* expression with the proliferative GC compartment and indicated distinct cellular expression patterns of the three major metacestode β-tubulin isoforms.

We next examined the local structural environment surrounding E198 in Tub-1, Tub-2, and Tub-3 *in silico* (Fig. 9E). Strikingly, the Y200-containing Tub-2 isoform displayed an E198-Y200 hydrogen bond, reproducing an interaction previously proposed in structural studies of Y200-containing, benzimidazole-refractory β-tubulins (Aguayo-Ortiz et al., 2013a,b; Garge et al., 2021). No corresponding interaction was observed in the F200-containing Tub-1 isoform. Tub-3, which likewise contains F200, displayed a distinct local configuration in which E198 formed a hydrogen bond with T165. Human β-tubulin class III (HsTBB3), which naturally contains Y200, was included as an additional reference and similarly displayed an E198–Y200 hydrogen bond. Thus, the presence of Y200 in Tub-2 is associated with a local interaction pattern around E198 resembling that previously implicated in reduced benzimidazole susceptibility.

Taken together, these findings identified Tub-2 as a β-tubulin isoform with two characteristics of particular relevance to the observed persistence of GC during ABZ treatment: its expression is strongly associated with the proliferative GC compartment, and its predicted structure displays an E198–Y200 interaction previously proposed to contribute to reduced benzimidazole susceptibility. These observations suggested that preferential expression of Tub-2 could contribute to the relative refractoriness of the GC compartment to ABZ. We therefore next sought to directly determine the cellular distribution of Tub-2 within the parasite.

### Tub-2 is associated with proliferating cells of the germinal layer

The transcriptomic analyses strongly associated *tub-2* expression with the proliferative GC compartment. We therefore sought to directly determine the cellular distribution of Tub-2 protein within the metacestode. A transcript-specific analysis by *in situ* hybridization was not feasible because *tub-1*, *tub-2*, and *tub-3* display a high degree of nucleotide sequence similarity, precluding the design of probes that could reliably discriminate *tub-2* transcripts from those of the other two predominantly expressed β-tubulin genes. We therefore employed the monoclonal Tub 2.1 antibody to specifically investigate the cellular distribution of Tub-2 protein.

Detailed epitope mapping previously localized the Tub 2.1 epitope to amino acids 431–436 within the variable C-terminal region of β-tubulin and demonstrated that recognition by this antibody is β-tubulin isotype-dependent rather than pan-specific (Yang et al., 2009). Sequence comparison of the major *E. multilocularis* β-tubulins showed that the corresponding C-terminal epitope is present in Tub-2 but differs in Tub-1 and Tub-3 (Fig. 8C). To experimentally verify the specificity of Tub 2.1 for *E. multilocularis* Tub-2, we heterologously expressed V5-tagged Tub-1, Tub-2, and Tub-3 in *E. coli*. Whereas all three recombinant proteins were detected by an anti-V5 antibody, Tub 2.1 exclusively recognized Tub-2, with no detectable cross-reactivity against Tub-1 or Tub-3 (Fig. S1A). Western blot analysis of parasite material further detected a Tub 2.1-reactive protein of the expected molecular size (50 kDa) in *E. multilocularis* lysates, with a particularly prominent signal in stem cell-enriched primary cell preparations (Fig. S1B). These experiments established Tub 2.1 as a suitable reagent for specifically monitoring Tub-2 protein in subsequent immunohistochemical analyses.

We combined Tub 2.1 immunostaining with a 5-h EdU pulse to relate Tub-2 expression to proliferative activity within the germinal layer. Tub 2.1 staining revealed a distinct population of Tub-2+ cells distributed throughout the germinal layer (Fig. 10A, B). Strikingly, virtually all cells undergoing DNA synthesis were Tub-2+: 99% of EdU+ cells also displayed Tub-2 immunoreactivity. Conversely, EdU+ cells represented 32 ± 5% of the total Tub-2+ cell population (1,432 cells analyzed). Thus, Tub-2 expression encompassed essentially the entire population of cells undergoing S phase during the 5-h labeling period but extended to a substantially larger population of cells that did not incorporate EdU during this interval.

Closer examination revealed distinct Tub-2 staining patterns among these populations (Fig. 10C). EdU+/Tub-2+ cells generally displayed a comparatively weak and more diffuse Tub-2 signal. In contrast, a subset of EdU-/Tub-2+ cells exhibited particularly intense Tub-2 immunoreactivity, frequently arranged in a characteristic spindle-shaped structure surrounding or adjacent to the nucleus. Thus, the absence of EdU incorporation in these cells was not associated with reduced Tub-2 abundance; instead, some of the most prominent Tub-2 structures were observed in cells that had not undergone detectable DNA synthesis during the preceding 5 h.

Together with the transcriptomic enrichment of *tub-2* under GC-rich conditions and its reduction following GC depletion, the near-complete overlap between EdU incorporation and Tub-2 immunoreactivity provided strong evidence for an association of Tub-2 with the proliferative GC compartment. Since EdU incorporation identifies cells undergoing DNA synthesis during the 5-h labeling period but does not encompass the entire GC population, we next sought to independently establish the identity of Tub-2+ cells using previously characterized molecular markers of *E. multilocularis* GC.

### Tub-2 colocalizes with established markers of *E. multilocularis* GC

To independently establish the identity of Tub-2+ cells, we examined the relationship between Tub-2 protein and previously characterized molecular markers of the *E. multilocularis* GC compartment. We first combined Tub2.1 immunostaining with WISH for *trim*, which has previously been established as a marker of GC in the metacestode (Koziol et al., 2015). Tub-2 immunoreactivity showed extensive overlap with 98% (± 2%) of *trim*-expressing cells (n = 463) being also Tub-2+ throughout the germinal layer (Fig. 11A), independently supporting the association of Tub-2 with the GC population.

We next examined *CIP2Ah*, which was identified as a marker associated with GC in our previous transcriptomic analyses (Herz et al., 2024). Combined staining for *CIP2Ah* and Tub-2 again revealed extensive overlap between the two markers in the metacestode germinal layer, with 97 ± 3% of *CIP2Ah*+ cells (n = 397) also being Tub-2+ (Fig. 11B). Colocalization of *CIP2Ah* and Tub-2 was also observed in developing brood capsules, where numerous cells expressing both markers were present (Fig. 11C). Thus, the association of Tub-2 with GC was independently supported by two previously established molecular markers and was observed both in the germinal layer of the metacestode vesicle and in developing brood capsules.

We additionally examined the spatial distribution of Tub-2+ cells within the germinal layer. Optical sections obtained immediately beneath the laminated layer predominantly revealed the superficial phalloidin-positive muscle network, whereas sections taken 8 µm deeper within the same confocal stack contained numerous Tub-2+ cells (Fig. 11D, E). Thus, Tub-2+ GC were predominantly located within deeper regions of the germinal layer rather than in the superficial muscle layer adjacent to the parasite surface.

At higher magnification, individual Tub-2+ cells again displayed the prominent spindle-shaped Tub-2 structures already observed in the EdU colocalization experiments (Fig. 11F). The morphology and intracellular position of these structures were suggestive of mitotic spindle formation. Together, the EdU colocalization, transcriptomic expression patterns, and independent colocalization with *trim* and *CIP2Ah* establish Tub-2 as a β-tubulin isoform strongly associated with the *E. multilocularis* GC compartment. The recurrent observation of spindle-like Tub-2 structures further raised the possibility that Tub-2 participates in mitotic spindle formation in dividing GC.

### Tub-2 is associated with proliferating cells in primary cell cultures and activated protoscoleces

Having established the association of Tub-2 with GC in the metacestode germinal layer, we next examined Tub-2 distribution in primary cell cultures and in the protoscolex, two additional developmental contexts containing proliferating GC. *E. multilocularis* primary cultures are initially strongly enriched in GC (Koziol et al., 2014) and have therefore been extensively used to investigate the biology of the parasite stem cell compartment (Koziol and Brehm, 2015). For technical reasons (2 day old aggregates are too small for WMIHF associated washing steps), immunostaining of primary cell aggregates was performed after 7 days of culture, when substantial cellular differentiation had already occurred. Nevertheless, the resulting aggregates contained large numbers of Tub-2+ cells distributed throughout the developing tissue (Fig. 12A). A three-dimensional reconstruction as well as individual confocal sections revealed abundant Tub-2+ cells within the aggregates.

We next investigated Tub-2 expression in activated protoscoleces. Following a 5-h EdU pulse, proliferating cells were particularly prominent in the posterior region of the protoscolex corresponding to the prospective neck region. Many of these EdU+ cells displayed Tub-2 immunoreactivity (Fig. 12B, C), consistent with the association between Tub-2 and proliferating cells observed in the metacestode germinal layer. Higher-magnification analysis of the posterior region confirmed the presence of individual EdU-positive cells associated with Tub-2 staining (Fig. 12C).

In addition to the signal associated with proliferating cells, however, activated protoscoleces displayed strong Tub2.1 immunoreactivity along the tegument, particularly in the anterior region encompassing the suckers and rostellum (Fig. 12B). Whether this signal reflects expression of Tub-2 in tegumental structures or recognition of an additional β-tubulin isoform carrying a compatible EGEF-containing Tub2.1 epitope remains unresolved. Consequently, although the association of Tub-2 staining with EdU+ cells was also evident in the protoscolex, the prominent additional tegumental signal limits the utility of Tub2.1 as a selective marker for GC in this developmental stage.

Together, these observations further supported the association of Tub-2 with proliferating cells across different developmental contexts of *E. multilocularis*. We therefore next addressed the conspicuous spindle-shaped Tub-2 structures repeatedly observed in the GC population and investigated whether Tub-2 participates in mitotic spindle formation.

### Tub-2 localizes to the mitotic spindle in dividing GC

The association of Tub-2 with the GC compartment raised the question of its functional role in these cells. In our previous analyses, particularly intense spindle-shaped Tub-2 structures were repeatedly observed in EdU-negative cells (Figs. 10C and 11F). Since EdU incorporation during a 5-h pulse identifies cells undergoing DNA synthesis but does not label cells that have subsequently progressed into mitosis, the morphology of these Tub-2 structures suggested that they might represent mitotic spindles. We therefore examined Tub-2 localization in mitotic cells using phosphorylation of histone H3 at serine 10 (pH3^S10^) as a marker of mitotic chromatin (Koziol et al., 2014).

Metacestode vesicles were simultaneously stained with the Tub2.1 antibody and an antibody against pH3^S10^. Within the germinal layer, pH3^S10^-positive cells consistently displayed prominent Tub-2 staining associated with the mitotic apparatus (Fig. 13A). At higher magnification, Tub-2 assumed a characteristic spindle-shaped distribution surrounding the pH3^S10^-positive condensed chromatin (Fig. 13B).

Analysis of individual mitotic cells in single confocal optical sections further resolved the spatial relationship between Tub-2 and mitotic chromatin. Tub-2-positive structures surrounded and extended around the condensed pH3^S10^-positive chromosomes in a pattern characteristic of the mitotic spindle (Fig. 13C, D). These observations identify Tub-2 as a component of the mitotic spindle in dividing *E. multilocularis* cells and explain the intense spindle-shaped Tub-2 staining observed in EdU-negative cells in the preceding experiments.

Together, these findings establish Tub-2 as a major β-tubulin isoform of proliferative GC and as a component of their mitotic spindle. We therefore finally asked how prolonged exposure to ABZ or ABZ-SO affects the Tub-2+ cell population in intact metacestodes.

### The Tub-2-positive cell population persists during prolonged ABZ treatment

Having established the association of Tub-2 with proliferative GC, we finally investigated whether prolonged benzimidazole treatment affected the abundance of the Tub-2+ cell population. Intact metacestode vesicles were treated for 10 days with 1 or 10 µM ABZ or ABZ-SO, followed by a 5-h EdU pulse. Vesicles were subsequently stained for EdU incorporation and Tub-2, allowing proliferative activity and the abundance of Tub-2+ cells to be assessed in parallel within the same treated metacestodes (Fig. 14A).

Consistent with our previous experiments, prolonged exposure to ABZ or ABZ-SO reduced GC proliferation. Quantification of EdU incorporation revealed a significant reduction in the proportion of EdU+ cells following treatment with either 1 or 10 µM ABZ and with either 1 or 10 µM ABZ-SO compared with the DMSO control (Fig. 14B). Importantly, however, EdU+ cells remained detectable under all treatment conditions.

In marked contrast to the reduction in proliferative activity, the proportion of Tub-2+ cells remained unchanged following treatment with either ABZ or ABZ-SO (Fig. 14C). Tub-2+ cells represented a similar fraction of the total cell population in control and benzimidazole-treated vesicles, with no significant differences detected between any treatment group and the DMSO control. Thus, although prolonged benzimidazole exposure reduced the fraction of GC undergoing DNA synthesis, it did not result in a detectable depletion of the Tub-2+ cell population.

Together with the persistence of proliferative and regenerative capacity demonstrated in the preceding experiments, these findings further support the conclusion that ABZ and ABZ-SO suppress GC proliferation without efficiently eliminating the Tub-2+ GC population. The persistence of these cells despite prolonged drug exposure is consistent with the relative refractoriness of the *E. multilocularis* stem cell compartment to ABZ and provides a potential cellular basis for the regenerative capacity retained following drug withdrawal.

## Discussion

Despite the long-standing use of benzimidazoles for the treatment of AE, the biological basis for their predominantly parasitostatic activity remains poorly understood. We therefore investigated how sustained ABZ exposure affects distinct cellular compartments of the *E. multilocularis* metacestode, with particular emphasis on GC that maintain parasite growth and regenerative capacity. The central finding of this study is that the pronounced effects of ABZ on the *E. multilocularis* metacestode are not accompanied by eradication of its regenerative GC compartment. This distinction provides, to our knowledge, the first direct cellular explanation for the predominantly parasitostatic rather than parasiticidal activity of benzimidazoles in AE. Previous clinical observations have demonstrated that parasite growth can resume after discontinuation of long-term benzimidazole treatment (Ammann et al., 1990; Ammann et al., 2015; Deibel et al., 2022), while experimental studies established GC as the only continuously proliferating somatic cell population of the metacestode and as the source of its extensive regenerative capacity (Spiliotis et al., 2008; Koziol et al., 2014). On this basis, we and others previously proposed that persistence of GC during benzimidazole treatment could account for parasite survival and subsequent regrowth (Schubert et al., 2014; Brehm and Koziol, 2014; Koziol and Brehm, 2015; Lundström-Stadelmann et al., 2019). The present data now provide direct experimental support for this model. GC remained detectable during prolonged ABZ exposure, and, critically, surviving cells retained developmental competence. Primary cells recovered from metacestodes after 14 days of ABZ treatment regenerated new vesicles after drug withdrawal, while GC-enriched primary cultures retained regenerative capacity even after 28 days of continuous exposure to 10 µM ABZ. Cells surviving this extended treatment were furthermore capable of generating parasite tissue following transplantation into a permissive host. Thus, persistence under ABZ does not merely represent residual metabolic activity or temporary cell-cycle arrest, but survival of a functionally competent cell population capable of re-establishing parasite tissue after drug withdrawal. This provides a plausible cellular basis for the clinically well-documented ability of *E. multilocularis* to persist during years of benzimidazole therapy and, in some patients, resume growth following treatment cessation (Ammann et al., 1990; Deibel et al., 2022; Autier et al., 2024; Deibel et al., 2025). Together, these findings support a model in which ABZ profoundly damages metacestode tissue while sparing a persistent regenerative cell compartment capable of re-establishing parasite tissue after drug withdrawal (Fig. 15A–C).

**Figure 15.**
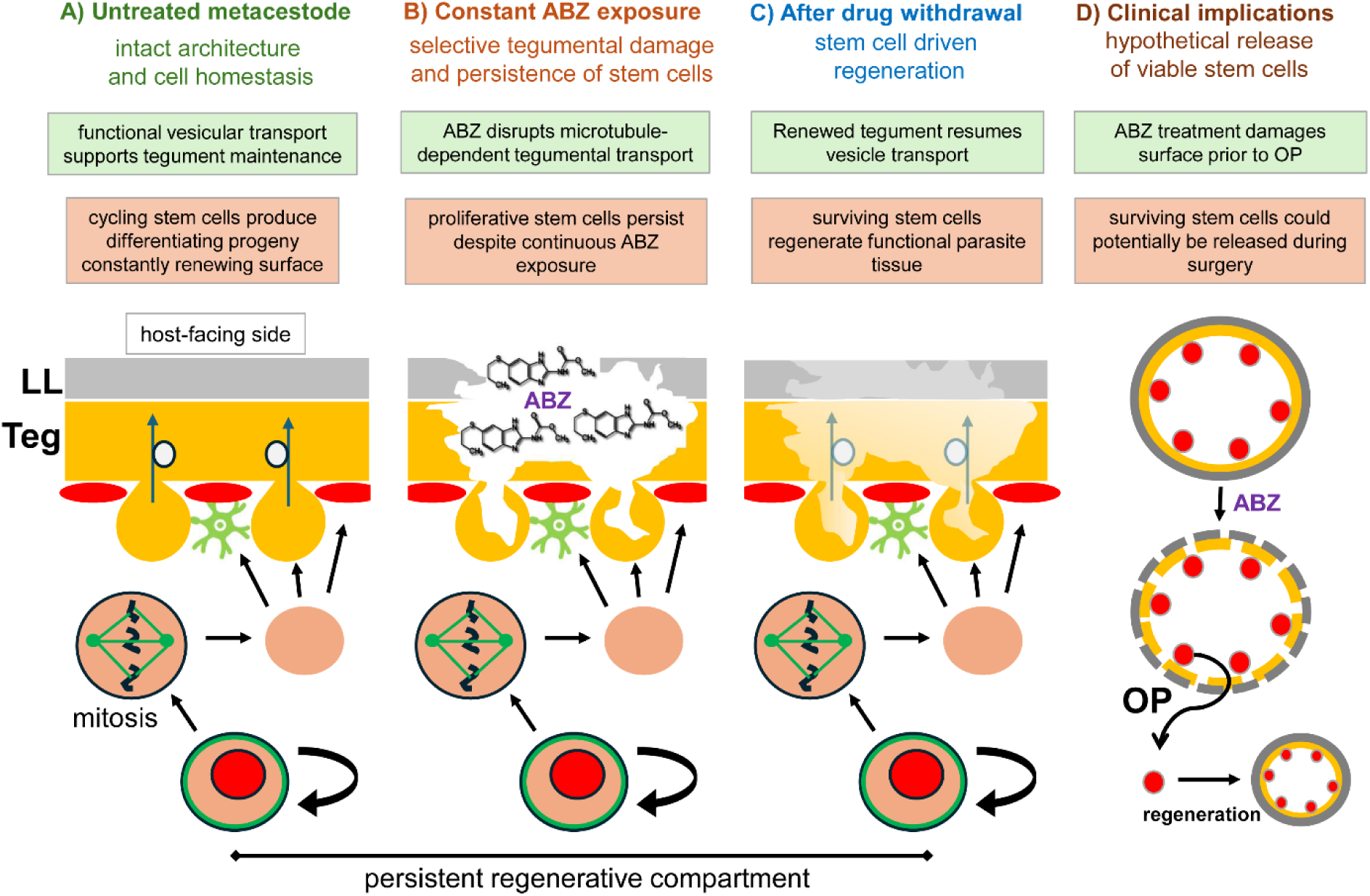
Proposed model for the cellular basis of the parasitostatic activity of albendazole against *E. multilocularis*. **(A)** In untreated metacestodes, continuous microtubule-dependent vesicular transport supports maintenance of the distal tegument, while proliferating germinative stem cells generate differentiating progeny required for tissue homeostasis and surface renewal. **(B)** During continuous ABZ exposure, disruption of microtubule-dependent tegumental transport results in progressive damage to the parasite surface and loss of metacestode integrity. In contrast, proliferative GC persist, maintaining a regenerative cell compartment despite prolonged drug exposure. **(C)** Following ABZ withdrawal, surviving GC retain developmental competence and can regenerate parasite tissue, including restoration of the tegument and its vesicular transport system. **(D)** Potential clinical implication of GC persistence during preoperative ABZ treatment. ABZ-induced damage to parasite architecture while viable regenerative cells remain present could hypothetically facilitate the release of such cells during surgical manipulation, followed by regeneration of parasite tissue. This scenario represents a hypothetical consequence of the cellular effects observed in this study and has not been demonstrated experimentally or clinically. LL, laminated layer; Teg, tegument; OP, surgical procedure.

Our findings further indicate that different cellular compartments of the metacestode differ markedly in their susceptibility to ABZ. One of the earliest detectable effects was the rapid appearance in the cytons of WGA-positive material associated with the tegument, preceding gross disruption of metacestode integrity. This is consistent with previous ultrastructural studies demonstrating pronounced tegumental alterations following benzimidazole treatment (Ingold et al., 1999; Küster et al., 2014) and with the recent observation in *M. corti* that tegumental microtubules are particularly sensitive to ABZ and ABZ-SO (Guarnaschelli and Koziol, 2025). In the latter study, benzimidazole treatment caused accumulation of secretory material within tegumental cytons and strongly reduced delivery of newly synthesized proteins to the distal tegument, directly linking disruption of microtubule function to impaired tegumental transport (Guarnaschelli and Koziol, 2025). The rapid accumulation of WGA+ cytons observed here is therefore compatible with inhibition of the continuous microtubule-dependent trafficking required to maintain the metabolically active parasite surface (Fig. 15A, B). In contrast, prolonged ABZ exposure did not result in a comparable depletion of differentiated muscle or neuronal cells. This difference does not necessarily imply that the β-tubulin isoforms expressed in these cells are intrinsically refractory to ABZ. Rather, the consequences of benzimidazole binding are likely to depend both on the molecular susceptibility of the β-tubulin isoform involved and on the extent to which a particular cellular process depends on dynamic microtubule function. Continuous long-range transport between tegumental cytons and the distal tegument may therefore be particularly vulnerable to interference with microtubule dynamics, whereas the survival of differentiated muscle or neuronal cells may be less acutely dependent on rapid microtubule turnover. Differential cellular responses to ABZ may thus reflect not only which β-tubulin isoforms are expressed, but also when and for which processes their microtubules are required.

The differential cellular susceptibility to ABZ may also have a molecular basis in the β-tubulin isoforms expressed by individual cell populations. Among the β-tubulin genes expressed in the metacestode, *tub-2* was by far the most abundant and showed a striking association with the GC compartment. This association was supported at the protein level by the extensive overlap of Tub-2 immunoreactivity with EdU incorporation and established GC markers, while its localization to mitotic spindles implicates Tub-2 in GC division. These observations are particularly noteworthy because Tub-2 differs from the other major metacestode β-tubulins at one of the best-established determinants of benzimidazole susceptibility: Tub-2 contains Y200, whereas Tub-1 and Tub-3 contain F200 (Brehm et al., 2000). Substitutions at positions 167, 198, and 200 are well-established determinants of benzimidazole susceptibility in parasitic nematodes, with F200Y representing one of the most frequently observed resistance-associated changes (Kwa et al., 1994; von Samson-Himmelstjerna et al., 2007; Furtado et al., 2014). Introduction of F200Y, as well as F167Y and E198A, into a benzimidazole-susceptible *Caenorhabditis elegans* background is sufficient to confer resistance, providing direct functional evidence for the contribution of these residues to benzimidazole susceptibility (Dilks et al., 2020).

Our structural analyses provide an additional link between the sequence characteristics of Tub-2 and its potential pharmacological properties. In Tub-2, Y200 formed a hydrogen bond with E198, reproducing an interaction previously proposed in structural models of Y200-containing, benzimidazole-refractory β-tubulins (Aguayo-Ortiz et al., 2013a, b). E198 is located within the benzimidazole-binding region, and alterations of the interaction network surrounding this residue have been proposed to affect the geometry and physicochemical properties of the binding site (Aguayo-Ortiz et al., 2013a, b). The E198–Y200 interaction observed in Tub-2 therefore provides a plausible structural basis for reduced ABZ susceptibility of this isoform. Together with the association of Tub-2 with GC, its localization to the mitotic spindle, and the persistence of Tub-2+ cells during prolonged ABZ exposure, these observations support a model in which Tub-2 contributes to the relative refractoriness of the regenerative cell compartment, as previously proposed on the basis of its expression pattern (Schubert et al., 2014; Brehm and Koziol, 2014; Koziol and Brehm, 2015). The present data establish an association rather than a causal relationship, however, and direct functional comparison of the individual β-tubulin isoforms will ultimately be required to test this model.

Importantly, this interpretation should not be reduced to a simple F200-sensitive/Y200-refractory dichotomy. The structural environment of the benzimidazole-binding region differs among Tub-1, Tub-2, and Tub-3 at several positions. Despite containing F200, Tub-3 displayed an alternative hydrogen bond between E198 and T165 in our structural model, indicating that the local configuration of E198 is not determined solely by residue 200. Structural and functional studies of β-tubulin resistance alleles similarly implicate several interacting residues within this region, including positions 165, 167, 198, and 200, in determining benzimidazole binding and susceptibility (Aguayo-Ortiz et al., 2013a, b; Dilks et al., 2020). Tub-1 and Tub-3 therefore cannot be assumed to represent uniformly ABZ-sensitive counterparts of Tub-2 simply because they contain F200, and direct biochemical or functional comparison will be required to establish the relative susceptibilities of the three isoforms.

The expression patterns of these isoforms nevertheless raise the possibility that β-tubulin composition contributes to the differential responses of parasite cell compartments. Whereas *tub-2* expression closely followed the abundance of GC, *tub-1* and *tub-3* showed markedly different expression profiles, being relatively enriched in conditions depleted of GC and dominated by differentiated cell populations. Although these transcriptomic data do not permit assignment of either isoform to a specific differentiated cell type, they are compatible with expression of Tub-1 and/or Tub-3 in differentiated compartments, potentially including the tegument. Such a distribution could contribute to the pronounced tegumental response to ABZ if one of these isoforms proves more ABZ-sensitive than Tub-2. Alternatively, or additionally, the strong tegumental phenotype may reflect the exceptional dependence of this compartment on continuous microtubule-mediated transport (Guarnaschelli and Koziol, 2025).

β-Tubulin sequence and cell-type-specific expression may represent only part of the explanation for GC persistence. The cellular consequences of benzimidazole binding are also likely to depend on when microtubule function becomes essential. GC may tolerate partial interference with microtubule dynamics during substantial portions of the cell cycle, whereas entry into mitosis creates an acute requirement for rapid microtubule assembly and reorganization to form a functional spindle. The localization of Tub-2 to mitotic spindles is therefore particularly relevant: if Tub-2 is intrinsically less susceptible to ABZ, preferential use of this isoform during mitosis could protect GC precisely at a stage at which inhibition of microtubule dynamics would otherwise be expected to have severe consequences. Conversely, the persistence of EdU+ cells during prolonged treatment indicates that ABZ does not simply impose a permanent block to cell-cycle progression. Thus, the response of an individual parasite cell to ABZ may ultimately be determined by an interplay between β-tubulin isoform structure, isoform expression, and the temporal requirement for dynamic microtubule function.

These findings also have implications for the development and evaluation of new drugs against AE. Because long-term parasite persistence ultimately depends on its regenerative GC compartment, pronounced morphological damage to metacestode vesicles or loss of metabolic activity alone cannot be considered sufficient indicators of parasiticidal efficacy. Compounds that efficiently disrupt the tegument or other differentiated structures may produce a strong phenotypic response while leaving developmentally competent GC intact. Future drug-screening strategies should therefore explicitly assess GC survival and proliferation and, importantly, determine whether regenerative capacity persists after drug withdrawal. Primary cell cultures, EdU-based analysis of GC proliferation, and regeneration assays provide complementary approaches to distinguish compounds that suppress metacestode function from those capable of eliminating the cells responsible for parasite persistence (Spiliotis et al., 2008; Schubert et al., 2014; Brehm and Koziol, 2014; Koziol and Brehm, 2015; Cheng et al., 2019). At the same time, β-tubulin should not be regarded as an exhausted target simply because current benzimidazoles are predominantly parasitostatic. Structurally distinct β-tubulin isoforms in different parasite cell compartments may provide opportunities for compounds capable of efficiently targeting the microtubule machinery required for GC division. The GC-associated Tub-2 isoform therefore represents a particularly relevant molecular target, and future screening of benzimidazole derivatives or other microtubule-directed compounds should include activity against Tub-2-associated GC rather than relying predominantly on damage to intact metacestode vesicles.

Our findings may also have implications for the perioperative use of benzimidazoles in AE. Radical, tumour-like resection remains the only established curative treatment for localized disease, with complete removal of viable parasite tissue representing the principal surgical objective (Brunetti et al., 2010; Hillenbrand et al., 2017; Grüner et al., 2017), and international recommendations have traditionally combined complete surgical removal with postoperative ABZ therapy, generally for at least two years (Brunetti et al., 2010). Recent clinical data suggest that postoperative treatment duration may be shortened in carefully selected patients following curative resection (Grüner et al., 2017; Calame et al., 2025; Deibel et al., 2025). Our results provide a possible biological rationale for the efficacy of postoperative ABZ even if GC themselves are relatively refractory to the drug. Residual GC would have to proliferate, generate differentiated progeny, and reconstruct functional metacestode tissue to establish a new lesion. Continuous ABZ exposure could interfere with this process, particularly with the establishment and maintenance of the tegument and its microtubule-dependent transport system (Ingold et al., 1999; Guarnaschelli and Koziol, 2025). Thus, postoperative ABZ need not necessarily eliminate every residual GC to prevent parasite regrowth; sustained interference with the generation and function of differentiated parasite tissue may be sufficient to suppress successful re-establishment of the metacestode. This is consistent with the clinical efficacy of prolonged benzimidazole treatment in controlling residual or unresectable AE despite its predominantly parasitostatic activity (Brunetti et al., 2010; Grüner et al., 2017; Autier et al., 2024; Deibel et al., 2025).

The situation may be different for ABZ administered before an otherwise curative surgical resection. Although international recommendations primarily emphasize radical resection followed by postoperative ABZ treatment (Brunetti et al., 2010), preoperative benzimidazole therapy is frequently used in contemporary surgical practice. In a German series, 22 of 33 patients (67%) received benzimidazole treatment before surgery, including ten treated for one to three months with explicitly neoadjuvant intent (Strohaeker et al., 2022). In another series, preoperative ABZ was administered to 16 of 23 patients (70%) undergoing laparoscopic and 51 of 70 patients (73%) undergoing open hepatic resection (Gloor et al., 2022), while a recent French-Swiss cohort reported preoperative ABZ in 111 of 195 patients (57%) undergoing hepatectomy (Rrupa et al., 2026). Preoperative treatment has also been deliberately employed as an inductive strategy preceding radical resection in individual patients (Bartels et al., 2020). Thus, preoperative benzimidazole exposure is common in AE surgery despite the absence of prospective evidence establishing a benefit in patients whose lesions are already considered directly resectable.

Our experimental findings raise a potential biological concern regarding this practice. ABZ rapidly impaired tegument-associated microtubule-dependent processes and subsequently compromised metacestode integrity, whereas proliferative and highly regenerative GC remained viable even during prolonged continuous exposure. Preoperative ABZ could therefore preferentially damage structural and differentiated parasite compartments while sparing precisely those cells with the greatest capacity to re-establish parasite tissue. Interestingly, histopathological analysis of human AE lesions has shown that ABZ treatment is associated with increased numbers of small Em2+ parasite particles within and around lesions, together with changes in the inflammatory response and parasite-associated structures (Ricken et al., 2017). Although the biological significance and cellular composition of these particles are unknown, this observation illustrates that ABZ can substantially alter the architecture of human AE lesions. In a patient scheduled for complete surgical removal, this creates a fundamentally different situation from postoperative therapy: the immediate objective is not long-term suppression of metacestode growth but physical removal of all viable parasite material without dissemination. Conceivably, surgical manipulation of an ABZ-damaged and structurally compromised lesion could facilitate the release of viable parasite cells into surrounding tissue or the operative field (Fig. 15D).

This possibility remains hypothetical, and there is currently no clinical evidence that preoperative ABZ increases postoperative recurrence. Indeed, a recent retrospective French-Swiss study identified the absence of preoperative ABZ as a factor associated with recurrence following hepatectomy (Rrupa et al., 2026). In that cohort, preoperative ABZ was associated with a lower risk of recurrence in multivariable analysis, arguing against a simple detrimental effect of preoperative treatment. However, only ten recurrences occurred among 195 patients, and the retrospective design cannot establish that preoperative ABZ itself caused the improved outcome or exclude contributions from patient selection, disease characteristics, surgical management, and subsequent therapy. Our data therefore do not indicate that preoperative ABZ is clinically harmful but identify a previously underappreciated biological consideration: ABZ can substantially affect parasite architecture while leaving the regenerative cell compartment intact. Given the widespread use of preoperative benzimidazole treatment and the absence of prospective evidence establishing its benefit in otherwise directly resectable AE, the biological and clinical consequences of this practice warrant specific evaluation. In particular, it will be important to determine whether preoperative treatment reduces the risk posed by viable parasite material during surgery or whether preservation of regenerative cells within structurally damaged lesions might, under some circumstances, counteract this presumed benefit.

Our study establishes that prolonged ABZ exposure can profoundly damage *E. multilocularis* metacestodes without eliminating the cells responsible for their regeneration. The persistence of proliferative GC that retain developmental competence after drug withdrawal provides a direct cellular explanation for a defining limitation of current benzimidazole chemotherapy: effective suppression of parasite growth without reliable parasite eradication. Our findings further identify differential β-tubulin isoform usage, and particularly the GC-associated Y200-containing Tub-2 isoform, as a plausible molecular component of this phenotype. Together, these observations shift the relevant endpoint for anti-echinococcal drug development from damage to the metacestode towards elimination of its regenerative potential. Future curative strategies should therefore be evaluated by their capacity to eradicate GC and prevent parasite regeneration, with the β-tubulin machinery of this cell compartment representing a particularly promising molecular target.

## Acknowledgements

This work was funded by the Manfred Wellhöfer foundation (Schwarzenbruck, Germany; grant 824000). The authors are indebted to Dirk Radloff for excellent technical assistance.

## Supplementary Material

**Table S1.**
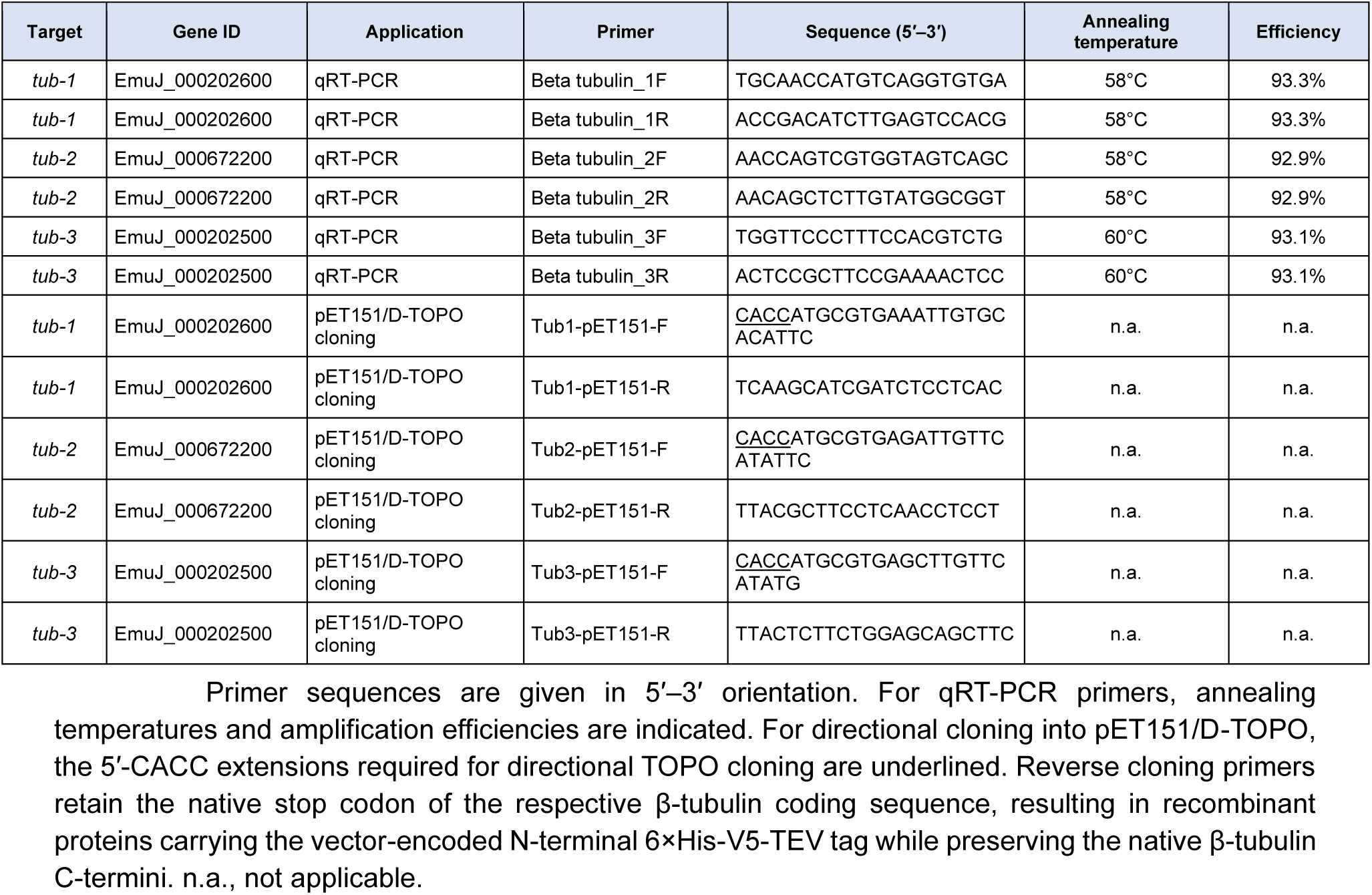
Oligonucleotide primers used in this study. Primer sequences are given in 5′–3′ orientation. For qRT-PCR primers, annealing temperatures and amplification efficiencies are indicated. For directional cloning into pET151/D-TOPO, the 5′-CACC extensions required for directional TOPO cloning are underlined. Reverse cloning primers retain the native stop codon of the respective β-tubulin coding sequence, resulting in recombinant proteins carrying the vector-encoded N-terminal 6×His-V5-TEV tag while preserving the native β-tubulin C-termini. n.a., not applicable.

**Figure S1.**
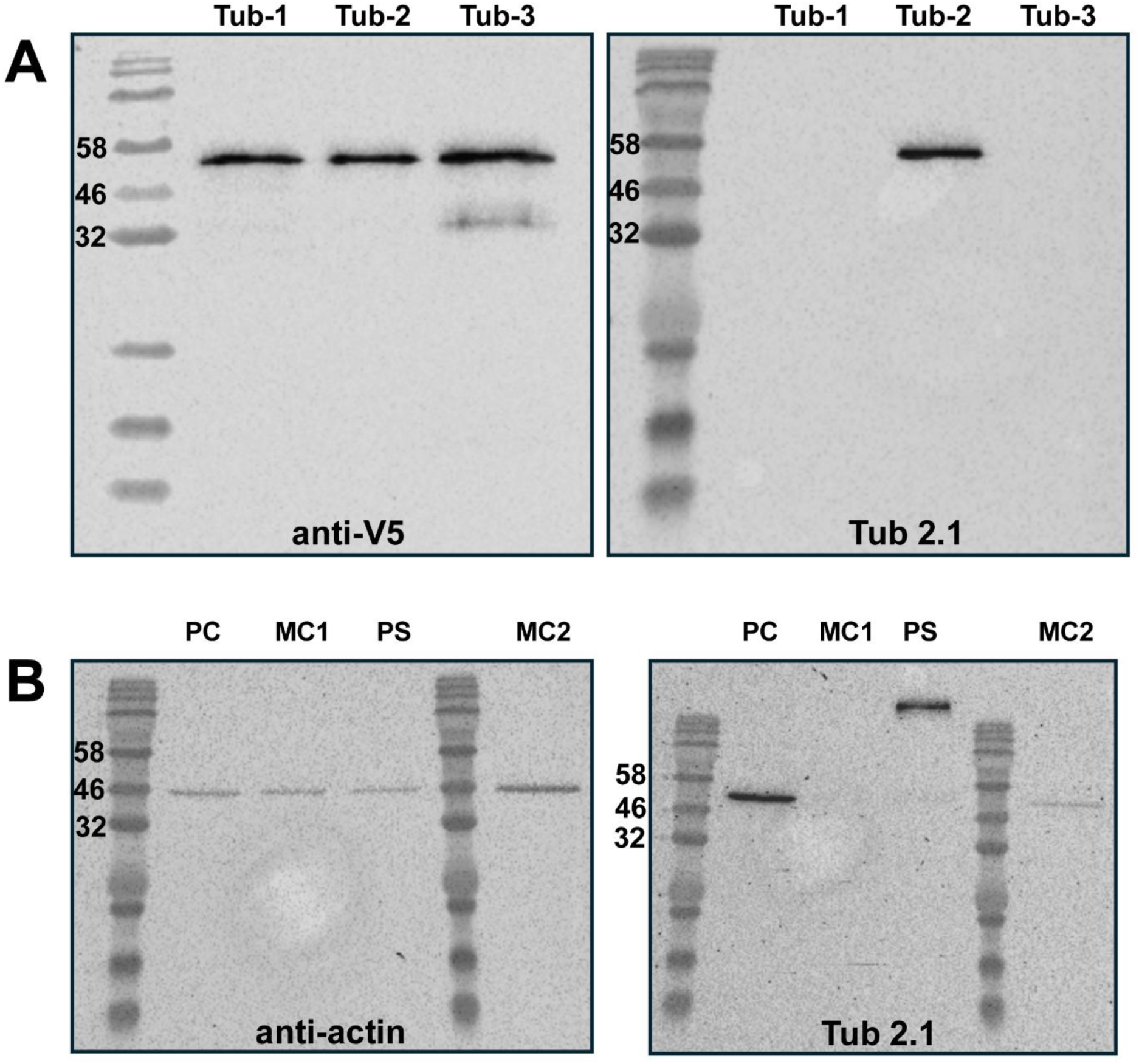
Validation of the Tub2.1 antibody for specific detection of *E. multilocularis* Tub-2. (A) Analysis of the specificity of the Tub2.1 antibody using heterologously expressed *E. multilocularis* β-tubulin isoforms. Tub-1, Tub-2, and Tub-3 were expressed as V5-tagged fusion proteins in *E. coli*. Purified recombinant proteins were separated by SDS-PAGE on 12.5% polyacrylamide gels and analyzed by Western blotting using an anti-V5 antibody (left) to detect the respective recombinant proteins or the Tub 2.1 antibody (right) to assess isoform specificity. Tub2.1 recognizes a C-terminal β-tubulin epitope containing the sequence EEEGEF, previously defined as its target epitope (Yang et al., 2009). (B) Detection of endogenous Tub-2 in different *E. multilocularis* preparations. Protein lysates from primary cells (PC), metacestode vesicles (MC1 and MC2), and protoscoleces (PS) were separated by SDS-PAGE on 12.5% polyacrylamide gels and analyzed by Western blotting using an anti-actin antibody as a control or the Tub2.1 antibody. For MC2, four times the amount of metacestode lysate used for MC1 was loaded. Protein marker sizes are indicated to the left of each blot.

